# Morpho-FM: spatial molecular reconstruction from routine H&E histology using transcriptomic foundation-model priors

**DOI:** 10.64898/2026.06.15.732498

**Authors:** Jinjin Huang, Xiao Feng, Lianghu Qu, Lingling Zheng

## Abstract

Routine haematoxylin and eosin (H&E) histology captures tissue architecture at clinical scale, but lacks a direct molecular readout of the transcriptional programmes that organise tumour epithelium, stroma, vasculature and immune compartments. Spatial transcriptomics provides this context, yet cost, workflow complexity and sparse sampling limit routine use. Most existing histology-to-expression models are trained de novo on small paired cohorts and therefore remain weakly constrained when extrapolating from sparse measurements to dense, tissue-wide molecular maps. Here we introduce Morpho-FM, a weakly supervised framework that predicts spatial gene expression from routine H&E whole-slide images by conditioning a pretrained single-cell transcriptomic foundation-model prior on local histological neighbourhoods. A lightweight morphology-to-transcriptome adapter maps cached whole-slide histology features into a transcriptomic decoder, enabling prediction at measured locations, dense full-section reconstruction, and re-aggregation to the original measurement support. Across harmonized prostate cancer benchmarks, Morpho-FM achieved the strongest overall performance among five representative methods, reaching mean per-gene Pearson correlations of 0.286 in rotating single-slide evaluation and 0.298 in multi-slide held-out validation. The framework reproduced this advantage across kidney cancer sections, achieved a mean correlation of 0.210 across 56 directed single-slide evaluations and retained measurable predictive signal after external transfer to clear-cell renal cell carcinoma sections. Controlled ablation analyses identified pretrained transcriptomic initialization as a reproducible source of performance gain exceeding that attributable to changes in the histology feature backbone. Beyond predictive accuracy benchmarks, Morpho-FM recovered *ERBB2*-enriched tumour compartments, boundary-associated molecular gradients, and annotation-aligned tissue domains across Xenium and HER2ST breast cancer datasets. Together, these results support transcriptomic foundation-model priors as an effective constraint for morphology-conditioned molecular decoding and demonstrate the potential of Morpho-FM to extend spatial transcriptomic insight across routine pathology sections.

## Introduction

Routine haematoxylin and eosin (H&E) histology remains the most widely deployed assay for tumour diagnosis and tissue assessment, because it captures cellular morphology, stromal organization and tissue architecture at whole-slide scale ^1^. Yet H&E images provide only an indirect view of the molecular programmes that shape tumour epithelium, stroma, vasculature and immune compartments ^2^. Transcriptional states that influence progression, therapy response and tissue microenvironmental organization often remain hidden behind morphologically similar regions. Bridging this gap between routine histology and spatially resolved molecular characterization has therefore become a central goal of computational pathology and spatial omics ^3–6^.

Spatial transcriptomics (ST) provides a direct route to this molecular context by measuring gene expression within native tissue architecture ^7–9^. These technologies have revealed spatial tumour states, tissue boundaries, immune niches and multicellular programmes that cannot be recovered from dissociated sequencing alone. However, their routine use remains limited by cost, workflow complexity, variable resolution and incomplete sampling of large tissue sections ^9,10^. In many practical settings, only sparse spatial measurements are available, whereas neighbouring or archived H&E sections are abundant. A scalable framework that can use sparse spatial transcriptomic supervision to molecularly reannotate routine histology would therefore expand the reach of spatial omics beyond directly assayed locations ^11^.

This need has motivated models that predict spatial gene expression from H&E images. Existing approaches have shown that histology contains measurable transcriptomic signal, but most are trained de novo on small paired cohorts and treat gene expression as a high-dimensional output vector to be learned almost entirely from local image supervision ^12–17^. This design leaves the output space weakly constrained, even though real expression profiles are structured by co-expression relationships, cell-state programmes and biological dependencies across genes. This limitation is particularly relevant in cancer, where local histomorphology reflects not only cell-intrinsic transcriptional states but also spatially contextual interactions within the tumour microenvironment that may not be adequately captured from limited paired examples alone.

Two classes of foundation model have advanced rapidly but remain largely disconnected in the task of inferring spatial gene expression from histology. In computational pathology, pretrained whole-slide encoders can represent tissue morphology at clinical scale ^18–20^. In single-cell genomics, transcriptomic foundation models have learned the structure of gene-expression space, including co-expression patterns, cell-state programmes and gene–gene dependencies ^21,22^. Existing histology-to-expression models have mainly asked whether stronger image representations can improve prediction from limited paired H&E-ST data. A more consequential question is whether morphology should be decoded through a transcriptomic foundation-model prior that constrains the space of biologically plausible expression states. In this view, pathology-derived representations identify local tissue context, while a single-cell transcriptomic prior restricts molecular decoding to states compatible with that context. We reasoned that coupling these two priors could address a central limitation of sparse paired datasets: learning not only where morphology varies across a tissue section, but also which transcriptional states, spatial distributions and tumour–microenvironment interactions those variations are likely to reflect.

Here we introduce Morpho-FM, a weakly supervised framework for spatial molecular reconstruction from routine H&E whole-slide images using transcriptomic foundation-model priors. Morpho-FM represents each measured spatial location as a local histological neighbourhood, maps cached whole-slide morphology features into the CellFM ^23^ transcriptomic latent space and decodes gene expression under sparse spatial transcriptomic supervision. The same trained model supports prediction at measured locations, dense full-section reconstruction and re-aggregation to the original measurement support for validation. Across prostate and kidney cancer benchmarks, external clear-cell renal cell carcinoma transfer, Xenium breast cancer reconstruction and HER2ST cross-platform analysis, Morpho-FM links benchmark accuracy to biologically interpretable molecular tissue architecture.

## Results

### Morpho-FM connects pathology-scale morphology to a transcriptomic foundation-model decoder

Morpho-FM was designed to test whether the pathology and transcriptomic foundation-model streams described above can be coupled for spatial molecular prediction. Rather than predicting each gene directly from an isolated H&E patch, the framework uses local histological neighbourhoods as the conditioning signal and decodes them through a pretrained single-cell transcriptomic foundation-model prior. This design separates the prediction problem into two coupled constraints: pathology-derived features capture where tissue architecture varies across the section, whereas the transcriptomic decoder constrains which expression states are plausible at those locations. To support both prediction at measured locations and dense full-section reconstruction, Morpho-FM implements this coupling within a shared coordinate system that links measured ST locations, cached whole-slide histology features, and dense prediction coordinates over the tissue mask (Fig. 1a).

**Fig 1.**
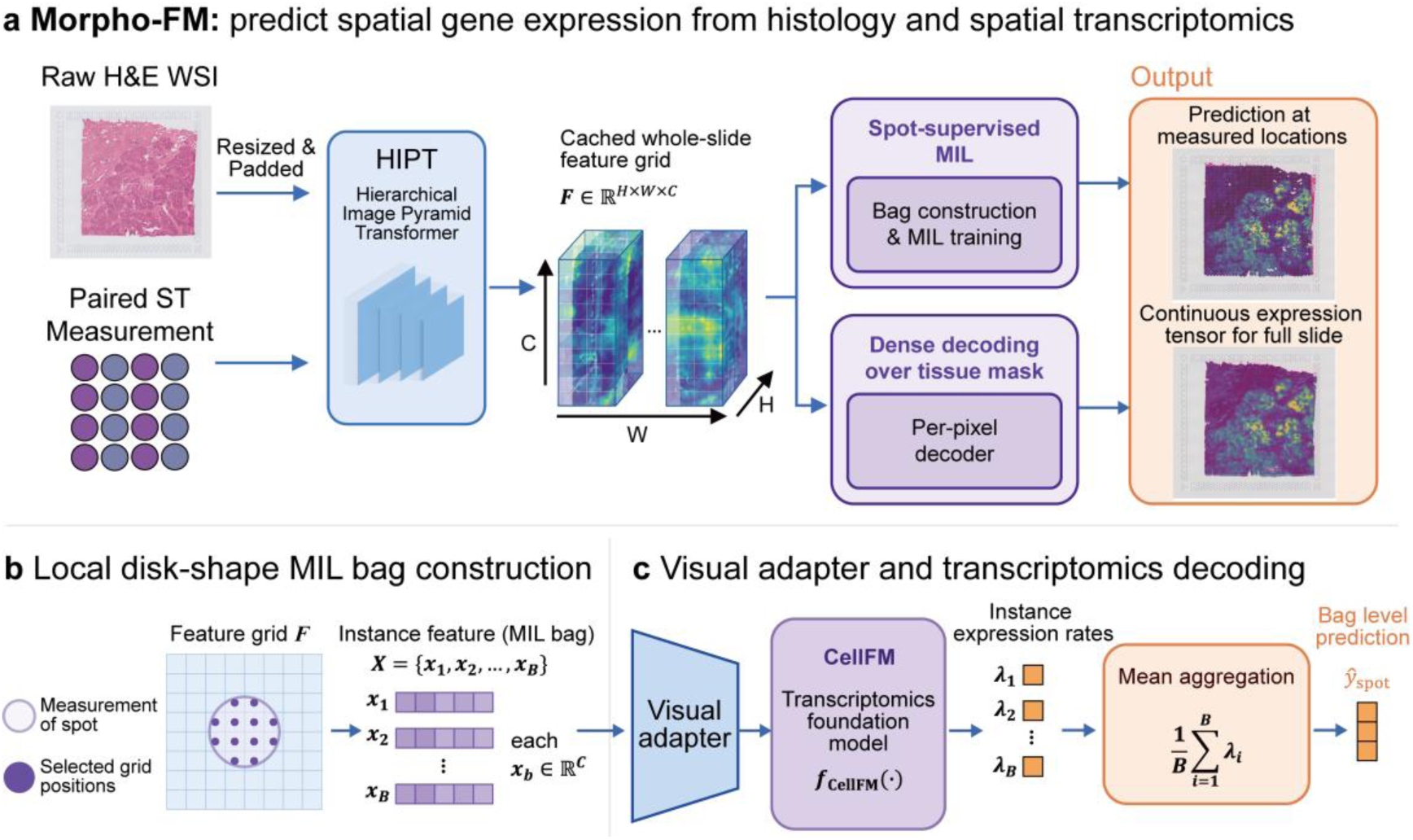
Morpho-FM couples local H&E morphology to transcriptomic foundation-model decoding in a shared coordinate system. a, Workflow from raw haematoxylin and eosin (H&E) whole-slide images and paired spatial transcriptomics (ST) measurements to two outputs: prediction at measured locations and dense full-section reconstruction. A cached whole-slide histology feature grid is constructed offline using a Hierarchical Image Pyramid Transformer (HIPT) encoder and used for both spot-supervised multiple-instance learning (MIL) and dense decoding over the tissue mask, with dense outputs re-aggregated to the original measurement support for consistency checking. **b,** Local disk-shaped MIL bag construction. For each ST measurement location, grid positions within the corresponding spot-radius neighbourhood are selected from the feature grid to form a MIL bag. **c,** Visual adapter and transcriptomic decoding. Instance features from the MIL bag are projected through a lightweight morphology-to-transcriptome adapter into CellFM, used here as the single-cell transcriptomic foundation-model decoder. The resulting instance-level expression rates are mean-aggregated to obtain the bag-level prediction.

The workflow begins by converting each raw H&E whole-slide image into cached whole-slide histology features using a Hierarchical Image Pyramid Transformer (HIPT) ^19^ encoder as the histology feature backbone (Fig. 1a). These features provide a reusable morphological representation for both spot-supervised training and dense decoding. Because measured assay locations and unmeasured tissue regions are indexed within the same spatial reference, Morpho-FM avoids constructing separate representations for spot-level prediction and tissue-wide reconstruction. Dense outputs can therefore be re-aggregated to the original measurement support, enabling quantitative consistency checking rather than serving only as qualitative super-resolution maps.

For sparse ST supervision, each measured location is represented by a local disk-shaped multiple-instance learning (MIL) bag sampled from the cached feature grid (Fig. 1b). Grid positions within the spot-radius neighbourhood are selected as instance features, so that each measurement is associated with its surrounding histological context rather than a single isolated image patch. This formulation matches the regional nature of ST measurements and provides the local H&E morphology used to condition transcriptomic decoding.

A lightweight morphology-to-transcriptome adapter then maps the instance features from each MIL bag into a transcriptomic decoder implemented with CellFM ^23^, a pretrained single-cell transcriptomic foundation model (Fig. 1c). The decoder produces instance-level expression rates, which are mean-aggregated to obtain a bag-level prediction aligned with the original ST measurement support. Thus, Morpho-FM does not simply append a regression head to a pathology encoder. Instead, it inserts a transcriptomic foundation model at the molecular readout stage, so that morphology-derived signals are interpreted within an expression space shaped by gene-gene dependencies and cell-state structure learned from large-scale single-cell data.

After spot-supervised training, Morpho-FM can be used in two complementary modes: prediction at measured locations and dense full-section reconstruction over the tissue mask (Fig. 1a). The resulting continuous expression fields can be visualized as tissue-wide molecular maps or re-aggregated to the original measurement support for comparison with measured ST data. We therefore organized the Results as a sequence of tests of the central hypothesis: first, whether this coupled decoder improves cross-section prediction; second, whether the gain is attributable to the transcriptomic foundation-model prior; and third, whether the resulting predictions and latent representations recover interpretable molecular tissue architecture beyond benchmark-level predictive performance.

### Morpho-FM improves cross-section histology-to-expression prediction in prostate cancer

We first evaluated Morpho-FM in prostate cancer, a setting designed to stress the limited-pairing regime in which transcriptomic priors should be most valuable. Four prostate cancer sections from the HEST-1k (HEST) collection ^24^ (INT25–INT28) were used, each comprising an H&E whole-slide image paired with spatial transcriptomics measurements. Five representative histology-to-expression methods were used for comparison: HisToGene, iStar, mclSTExp, sCellST and HiST. To ensure that performance differences reflected modelling rather than data handling, all methods were evaluated using the same preprocessing and data-splitting pipeline, with spatial coordinates aligned to the H&E image space and predictions assessed on a common prostate gene panel (Supplementary Note 1).

To test performance under the most data-limited condition, each prostate section was used in turn as the sole training slide, with each remaining section held out for testing, yielding 12 ordered train–test comparisons. Across these comparisons, Morpho-FM showed the strongest overall performance, achieving the highest mean per-gene Pearson correlation (mean, 0.286; median, 0.241). This exceeded the best-performing baseline, iStar (mean, 0.155; median, 0.132), as well as the other methods (mean, 0.038–0.055; Fig. 2a,b and Supplementary Figs. 1–3). The improvement was consistent across splits: Morpho-FM ranked first in all train–test directions. Performance was highest for INT27-to-INT28 (mean Pearson correlation, 0.367), whereas even the lowest-scoring Morpho-FM direction, INT25-to-INT26, outperformed iStar on the same test section (0.215 versus 0.149). The largest separation was observed for INT28-to-INT26 (0.223 versus 0.068; Supplementary Table 1).

**Fig 2.**
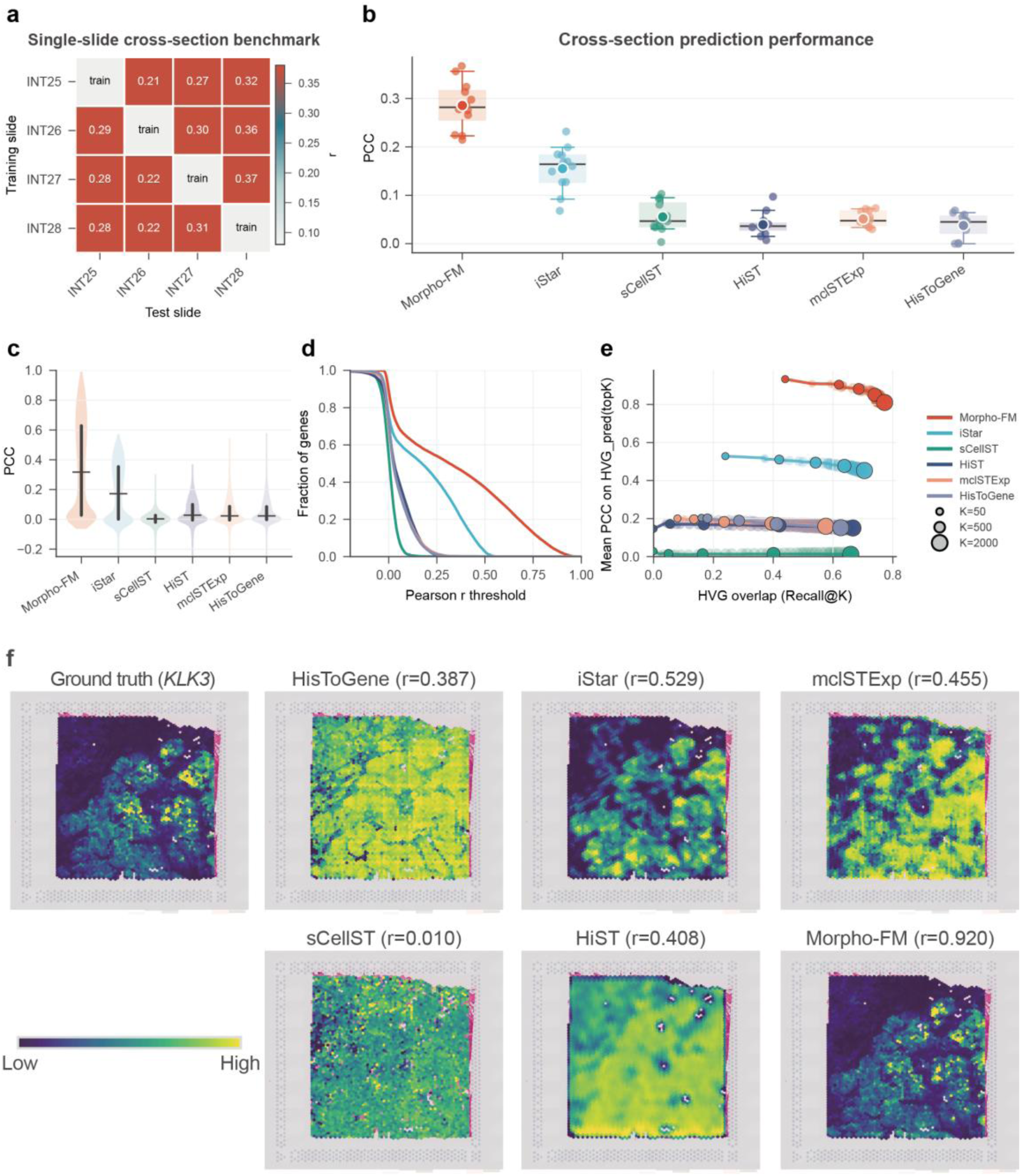
Single-slide cross-section benchmarking of Morpho-FM on prostate spatial transcriptomics data. a, Rotating single-slide train-test design across four prostate cancer sections, yielding 12 directed train-test comparisons. Heatmap values indicate the Morpho-FM mean per-gene Pearson correlation for each train–test direction; diagonal entries denote the training slide. **b,** Mean per-gene Pearson correlation across all 12 train-test directions for Morpho-FM and competing methods. Points denote individual train-test directions, and boxplots summarize the distribution across directions. **c,** Per-gene Pearson correlation distributions for the representative INT25-to-INT28 transfer comparison. **d,** Fraction of genes with Pearson correlation above each threshold in the INT25-to-INT28 comparison. **e,** Highly variable gene (HVG) recovery and prediction quality across top-*K* predicted gene sets. Recall@*K* measures overlap with measured HVGs, the x axis shows the mean Pearson correlation for the predicted top-*K* gene set, and point size denotes *K* = 50, 500 or 2,000. **f,** Spatial visualization of *KLK3* expression in the representative INT25-to-INT28 comparison. Ground truth and method predictions are shown after re-aggregation to the original measurement support; values in parentheses denote Pearson correlation coefficients between predicted and measured *KLK3* expression.

Morpho-FM’s improvement extended across the gene panel rather than being restricted to a small subset of genes. In the representative INT25-to-INT28 comparison, Morpho-FM shifted the distribution of per-gene correlations towards higher values and preserved the largest proportion of genes above increasingly stringent Pearson correlation thresholds (Fig. 2c,d). Similar trends were observed across the remaining train–test directions (Supplementary Figs. 1 and 2). We next tested whether this advantage persisted among highly variable genes (HVGs), which capture prominent spatial expression variability across measurement locations ^25–27^. Across top-*K* HVG cutoffs, Morpho-FM consistently occupied the high-recovery, high-correlation regime, recovering a larger fraction of measured HVGs while maintaining a higher mean Pearson correlation on the predicted HVG set than any baseline method (Fig. 2e and Supplementary Fig. 3). For the top 1,000 HVGs, Morpho-FM achieved a mean per-gene Pearson correlation of 0.778, compared with 0.443 for iStar and lower values for the remaining methods.

To determine whether improved gene-level accuracy translated into spatially coherent marker reconstruction, we examined *KLK3*, which encodes prostate-specific antigen (PSA) and is a clinically relevant marker in prostate adenocarcinoma^28^. In the representative INT25-to-INT28 comparison, Morpho-FM recapitulated the main spatial structure of the observed *KLK3* expression pattern after re-aggregation to the original measurement support (Fig. 2f). The reconstructed map captured broad low-expression regions and focal high-expression hotspots, yielding a Pearson correlation of 0.920 at the re-aggregated measurement locations (Fig. 2f). This marker-level result linked the genome-wide benchmark advantage to spatial reconstruction at measured locations.

We next asked whether the same advantage persisted when more paired sections were available. In a rotating multi-slide setting, two prostate sections were used for training, a third for validation and the remaining section for held-out testing. Across four held-out folds, Morpho-FM again ranked first among all methods, achieving a mean per-gene Pearson correlation of 0.298 (median, 0.260; top-1,000 HVG mean, 0.785; Extended Data Fig. 1a,b and Supplementary Fig. 4). The strongest baseline was again iStar (mean, 0.114; median, 0.092; top-1,000 HVG mean, 0.315), whereas the remaining baselines achieved mean Pearson correlations of 0.052-0.066 (Extended Data Fig. 1b). Morpho-FM was the top-performing method for each held-out prostate section, with fold-wise mean Pearson correlations ranging from 0.235 to 0.369 (Extended Data Fig. 1a). Together, the single-slide and multi-slide prostate benchmarks show that the coupled morphology-to-transcriptome decoder improves cross-section prediction under both sparse and richer supervision, rather than only under one favourable training regime.

### Morpho-FM generalizes histology-to-expression prediction to kidney cancer and external clear-cell renal cell carcinoma sections

We next tested whether the prostate result reflected a generalizable modelling advantage or a tissue-specific fit. Applying the same rotating single-slide design to eight HEST kidney cancer sections ^29^ yielded 56 directed train-test comparisons. Morpho-FM again achieved the highest mean per-gene Pearson correlation across directions (mean, 0.210; median, 0.194; top-1,000 HVG mean, 0.570), outperforming iStar, the strongest baseline (mean, 0.049; median, 0.038; top-1,000 HVG mean, 0.154), and all remaining methods, which had mean Pearson correlations below 0.027 (Fig. 3a and Supplementary Table 2). As in prostate cancer, Morpho-FM ranked first across the rotating comparisons, indicating that its advantage was reproduced in a second tumour context with different tissue architecture and expression programmes.

**Fig 3.**
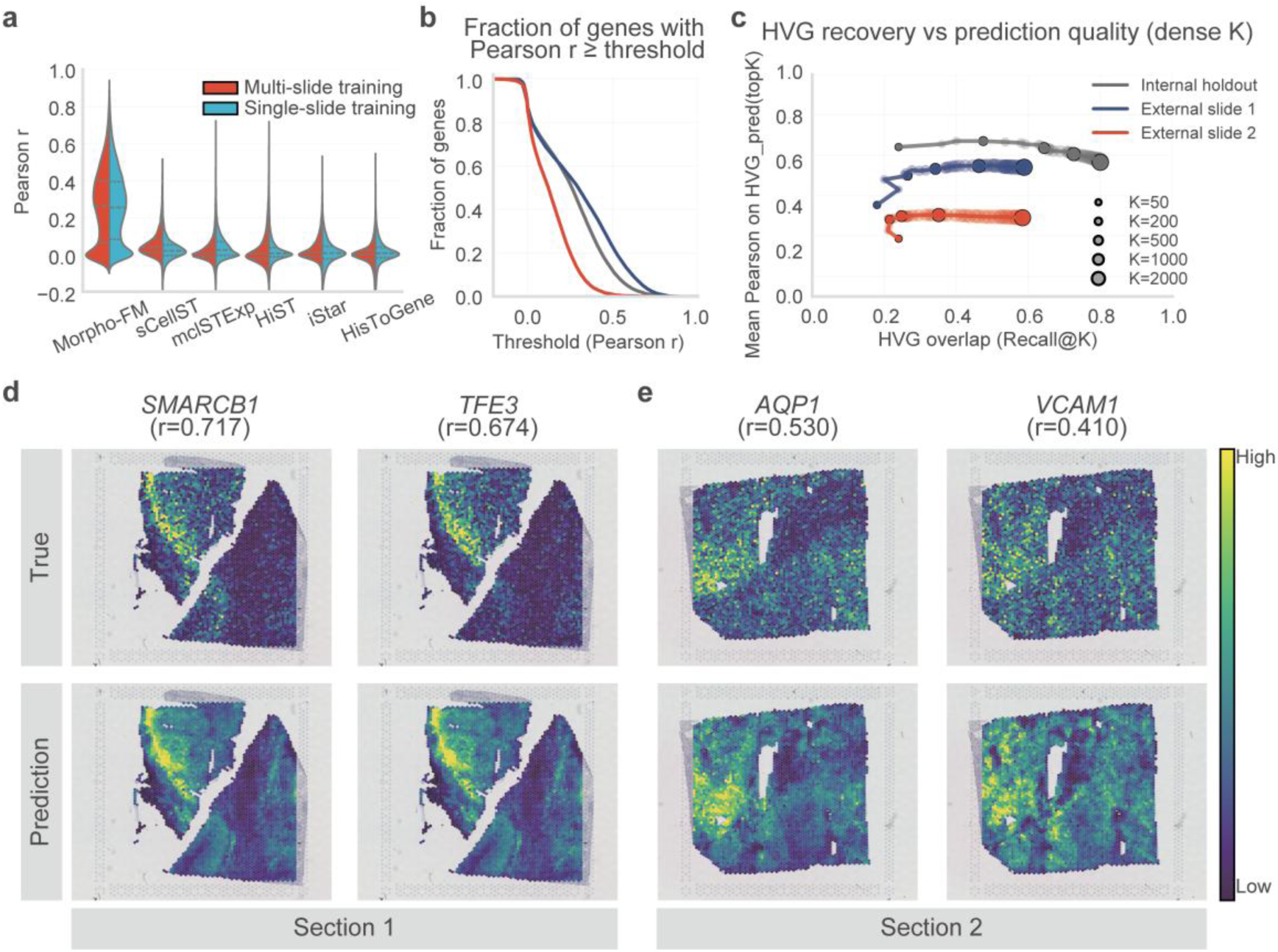
Morpho-FM reproduces kidney benchmark performance and transfers to external clear-cell renal cell carcinoma sections. a, Split violin plots of per-gene Pearson correlations across methods. Multi-slide and single-slide training results are shown on the left and right halves of each violin, respectively. **b,** Fraction of genes exceeding Pearson correlation thresholds for the internal kidney holdout and two external clear-cell renal cell carcinoma (ccRCC) sections. **c,** HVG recovery versus prediction quality across top-*K* cutoffs; point size denotes K. **d,e,** Representative external-section spatial maps comparing measured expression (top) and Morpho-FM prediction (bottom). In external section 1, *SMARCB1* and *TFE3* were recovered with *r* = 0.717 and *r* = 0.674, respectively (**d**). In external section 2, *AQP1* and *VCAM1* were recovered with *r* = 0.530 and *r* = 0.410, respectively (**e**). Colour scale indicates relative expression within each gene map.

We then moved from internal cross-section benchmarking to external transfer. A kidney-trained Morpho-FM model was selected using six HEST kidney sections for training, INT24 for validation and INT21 for held-out internal testing. On INT21, Morpho-FM achieved a mean per-gene Pearson correlation of 0.255 (median, 0.261; top-1,000 HVG mean, 0.620), remaining above the baseline methods on the same held-out kidney section (Fig. 3a). The resulting checkpoint was then applied without additional fitting, target-section-specific tuning or hyperparameter adjustment to two external clear-cell renal cell carcinoma (ccRCC) Visium sections ^30^. Morpho-FM retained measurable predictive signal in both external sections. In external sections 1 and 2, respectively, it achieved mean per-gene Pearson correlations of 0.294 and 0.138, section-level global expression correlations of 0.504 and 0.467, and mean spot-level expression-profile Pearson correlations of 0.658 and 0.501 (Fig. 3b,c). The reduction in external section 2 illustrates the difficulty of cross-dataset transfer, but the retained global and spot-level correlations show that the model did not collapse when moved outside the benchmark source cohort.

Representative marker maps showed that external predictions retained biological interpretability beyond numerical correlation alone. In the higher-performing external section 1, Morpho-FM recovered the spatial patterns of *SMARCB1* (*r* = 0.717), a SWI/SNF-complex component, and *TFE3* (*r* = 0.674), which is associated with MiT/TFE-family transcriptional programmes in renal tumours ^31,32^. In external section 2, the model recapitulated the distributions of *AQP1* (*r* = 0.530) and *VCAM1* (*r* = 0.410), markers of proximal tubular differentiation and inflammatory adhesion signalling, respectively ^33,34^ (Fig. 3d,e). Additional marker-gene comparisons in both external sections showed similar recovery of spatial patterns associated with renal tumours (Supplementary Fig. 10). Together, these kidney analyses extend the prostate benchmark by showing both reproducible method ranking in a second tumour type and preservation of interpretable molecular signal after transfer to independently processed ccRCC sections.

### Pretrained CellFM initialization is the dominant controlled source of Morpho-FM performance gain

The benchmark results were consistent with the hypothesis that a transcriptomic foundation-model decoder improves morphology-conditioned expression prediction, but benchmark comparisons alone do not identify the source of this advantage. We therefore isolated the contribution of pretrained transcriptomic initialization by comparing Morpho-FM models initialized with either pretrained CellFM weights or randomly initialized CellFM weights under matched multi-slide and single-slide prostate training settings. All remaining implementation details were controlled as described in Supplementary Note 2, enabling a direct test of whether pretrained single-cell transcriptomic priors improve prediction when training data, model architecture and optimization settings are otherwise held fixed.

Pretrained CellFM initialization improved prediction in both matched training settings, as measured by mean per-gene Pearson correlation and supported by per-gene correlation distributions and Pearson-threshold curves (Fig. 4a,b). In the multi-slide setting, the pretrained-initialized Morpho-FM model achieved a mean per-gene Pearson correlation of 0.360, compared with 0.299 for random initialization, an absolute increase of 0.061. In the single-slide setting, the corresponding values were 0.346 and 0.265, an absolute increase of 0.081. The larger gain in the single-slide comparison indicates that pretrained transcriptomic structure is especially beneficial when paired spatial transcriptomic supervision is sparse, supporting its role as a constraint on morphology-conditioned expression prediction.

**Fig 4.**
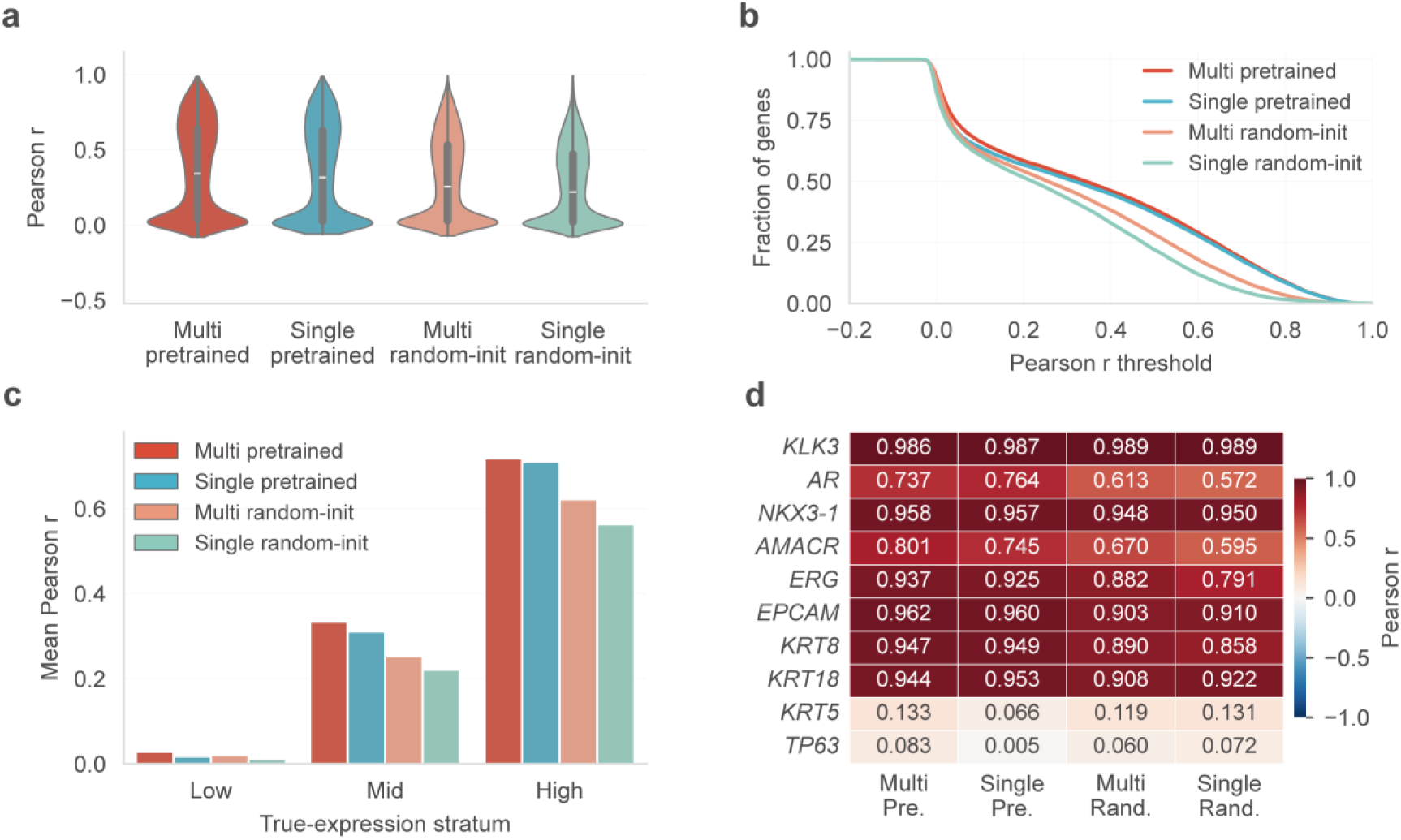
Pretrained CellFM initialization improves Morpho-FM performance across prostate training settings. a, Per-gene Pearson correlation distributions for Morpho-FM models with pretrained or random CellFM initialization under matched multi-slide and single-slide prostate training settings. **b,** Fraction of genes with Pearson correlation above each threshold for the four model variants shown in **a**. **c,** Mean per-gene Pearson correlation stratified by measured gene-expression level. Genes were grouped into low-, mid- and high-expression strata based on their measured expression levels, and performance is shown for each model variant. **d,** Gene-level Pearson correlations for a fixed prostate marker panel across the four model variants. Heatmap values indicate Pearson correlation coefficients between predicted and measured expression for each marker gene.

Expression-stratified analysis showed that the gain from pretrained transcriptomic initialization was not a uniform offset across all genes. Gains were small among low-expression genes, for which per-gene correlations remained low across the pretrained and randomly initialized models in both training settings, but increased in the mid- and high-expression strata (Fig. 4c). This pattern was reproduced in both the multi-slide and single-slide settings, with the largest improvement observed for high-expression genes in the data-limited single-slide comparison. A fixed prostate marker panel provided a gene-level view of the same effect: pretrained initialization improved correlations for several markers, including *AR*, *AMACR*, *ERG*, *EPCAM* and *KRT8*, whereas *KLK3* showed near-ceiling correlations across all model variants (Pearson r ≈ 0.99; Fig. 4d).

Thus, the gain from pretrained transcriptomic initialization was not driven by a single marker, but was most evident among genes with detectable spatial expression structure. We next asked whether a comparable gain could be obtained by changing the histology feature backbone while keeping the pretrained transcriptomic decoder fixed. Replacing HIPT features with ImageNet-pretrained ResNet-50 features produced comparable mean per-gene Pearson correlations in both prostate settings: ResNet-50 was slightly higher in the single-slide comparison, whereas HIPT was slightly higher in the multi-slide comparison (Supplementary Fig. 11). Expression-stratified summaries, Pearson-threshold curves and marker-panel correlations similarly showed no stable advantage for either histology feature backbone.

These controls do not indicate that histology features are unimportant; local morphology remains the conditioning signal for all Morpho-FM predictions. Rather, they separate two possible explanations for the benchmark gains. Within the present Morpho-FM design, replacing the histology feature backbone did not provide a consistent additional gain, whereas pretrained transcriptomic initialization produced a reproducible improvement in both data-rich and data-limited settings. The ablations therefore support the central design claim: in sparse H&E-ST learning, constraining the molecular decoder with a transcriptomic foundation-model prior is a clearer and more reproducible source of improvement than simply changing the upstream image feature backbone.

### Morpho-FM enables measurement-constrained reconstruction of molecular tissue architecture in Xenium breast cancer

Having established robust benchmark performance and isolated the contribution of transcriptomic initialization, we next asked whether Morpho-FM could do more than predict expression at measured coordinates. This task reflects the intended use case of molecularly reannotating tissue architecture, in which sparse spatial transcriptomic measurements are used to project molecular information across H&E-stained sections while retaining quantitative agreement with measured coordinates. Xenium provides a stringent setting for this analysis because its cell-resolved measurements support evaluation both at measured Xenium locations and after re-aggregation of dense predictions to the original measurement support ^35^. We applied Morpho-FM to two consecutive 10x Genomics Xenium formalin-fixed paraffin-embedded (FFPE) breast cancer sections ^36^, denoted section 1 (S1) and section 2 (S2). After matching the measured gene panel to the CellFM vocabulary, 306 genes were retained for downstream analysis. We evaluated four train–test settings: S1-to-S1, S1-to-S2, S2-to-S2 and S2-to-S1 (Fig. 5a).

**Fig 5.**
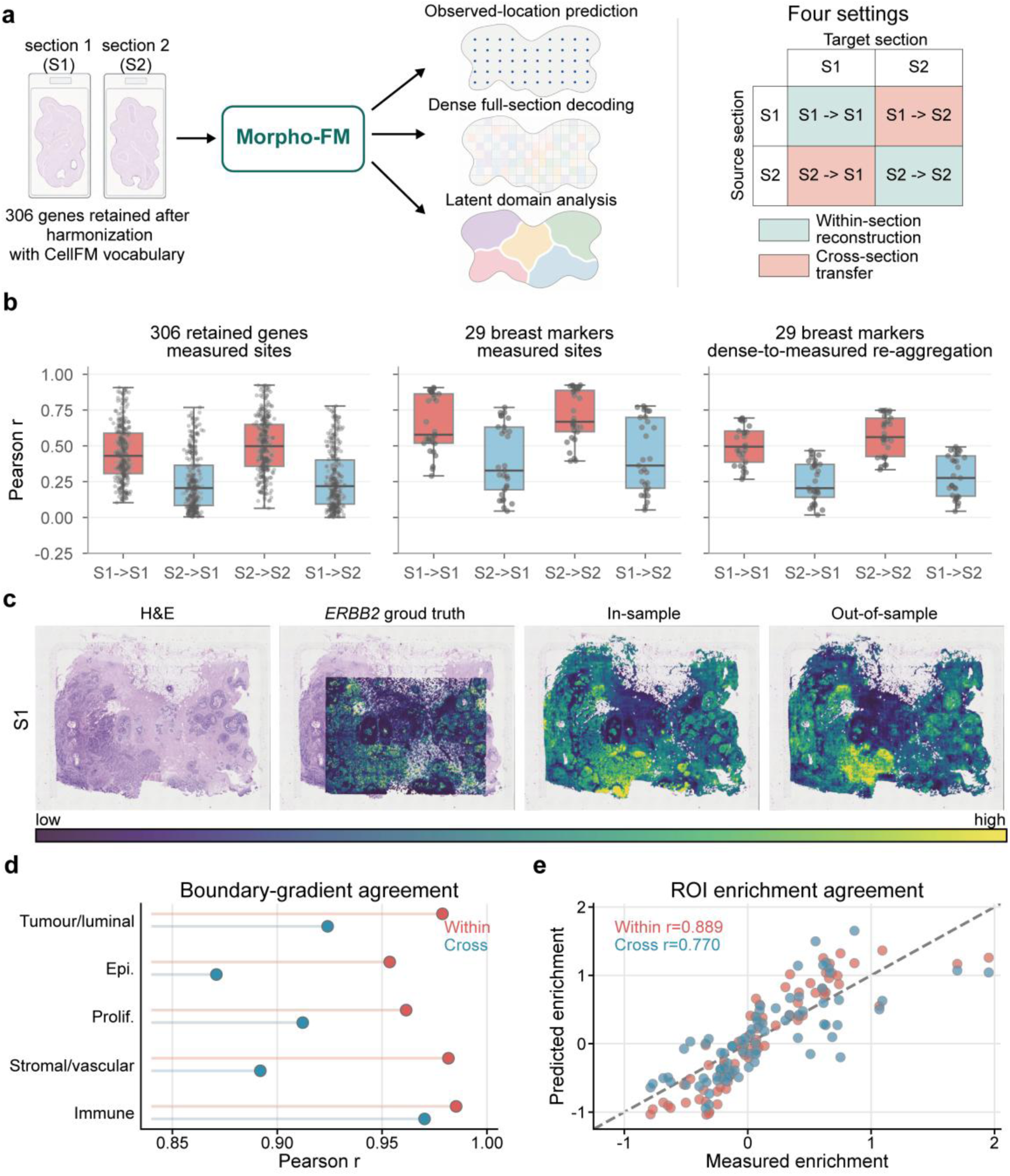
Measurement-constrained dense reconstruction in Xenium breast cancer sections. a, Schematic of the Xenium reconstruction analysis. Two consecutive Xenium formalin-fixed paraffin-embedded (FFPE) breast cancer sections, section 1 (S1) and section 2 (S2), were harmonized with the CellFM vocabulary, yielding 306 retained genes. Morpho-FM was evaluated for prediction at observed Xenium measurement locations and dense full-section reconstruction across four reciprocal settings: S1-to-S1, S1-to-S2, S2-to-S2 and S2-to-S1. **b,** Prediction performance across the four settings for all 306 retained genes, a fixed 29-gene breast marker panel, and the same marker panel after re-aggregation to the original Xenium measurement support. Points denote individual genes; boxplots summarize per-gene Pearson correlation distributions. **c,** Representative *ERBB2* reconstruction in S1, showing the haematoxylin and eosin (H&E) image, measured Xenium expression, within-section prediction and cross-section prediction. Colour intensity indicates relative expression. **d,** Boundary-gradient agreement between measured and predicted signature profiles across tumour-adjacent spatial bins. **e,** Region of interest (ROI) enrichment agreement between measured and predicted Xenium signatures across curated tissue regions. Each point denotes an ROI-signature pair.

Across these settings, within-section reconstruction consistently outperformed cross-section transfer, as expected, but reciprocal transfer retained measurable signal for biologically structured marker sets. For all 306 genes, mean per-gene Pearson correlations at measured Xenium locations were 0.448 and 0.501 for within-section prediction in S1 and S2, respectively, and decreased to 0.249 and 0.266 under reciprocal cross-section transfer (Fig. 5b and Supplementary Table 3). A fixed 29-gene panel spanning tumour/luminal, epithelial, proliferation, stromal, vascular and immune programmes showed higher absolute correlations while preserving the same hierarchy, with correlations of 0.651 and 0.708 within section compared with 0.396 and 0.427 across sections. Dense grid-level predictions showed a similar pattern after re-aggregation to the original Xenium measurement support, with mean Pearson correlations of 0.503 and 0.571 within section and 0.247 and 0.287 across sections (Fig. 5b and Supplementary Table 4). Thus, dense full-section reconstruction remained quantitatively coupled to the measured support, rather than functioning only as a visual extrapolation. Marker-category summaries further indicated that epithelial markers showed the strongest reproducibility, whereas vascular and immune markers showed larger cross-section loss (Supplementary Fig. 12). Representative marker-class maps for *FOXA1*, *EPCAM*, *MKI67*, *LUM*, *PECAM1* and *PTPRC* further illustrate these trends at both measured and dense supports (Extended Data Fig. 3).

*ERBB2* provided a marker-level example of how prediction at measured locations related to dense full-section reconstruction. In both Xenium sections, Morpho-FM recovered the major *ERBB2*-high compartments observed in the measured Xenium counts and preserved the broad contrast between tumour-dominant high-expression regions and surrounding low-expression tissue (Fig. 5c). Observation-level *ERBB2* correlations were high in the within-section setting, reaching 0.907 in S1 and 0.924 in S2, and remained positive under reciprocal transfer, with correlations of 0.778 for S1-to-S2 and 0.769 for S2-to-S1. Re-aggregation of dense predictions to the original measurement support produced lower, but still structured, correlations: 0.565 and 0.692 within section and 0.397 and 0.392 across sections. Dense decoding extended the *ERBB2* expression field from measured Xenium locations to 392,884 tissue-valid grid locations in S1 and 355,487 in S2, providing a tissue-wide molecular view of *ERBB2*-rich tumour compartments anchored to the same measurement support used for quantitative evaluation. Together with the representative marker-class examples in Extended Data Fig. 3, this result shows that dense reconstruction extended across tumour/luminal, epithelial, proliferation, stromal, vascular and immune marker programmes rather than being limited to *ERBB2*.

We further asked whether dense predictions preserved molecular organization at tissue boundaries and regions of interest (ROIs), where histological architecture and expression programmes are spatially coupled. Using curated tissue-compartment annotations, we computed the signed distance to the tumour edge and compared measured and predicted signature profiles across tumour-adjacent spatial bins (Fig. 5d). Within-section dense profiles closely matched measured boundary trends, with signature-wise Pearson correlations of 0.954-0.985. Cross-section profiles preserved the same broad gradients, although with lower correlations of 0.871-0.970. At the ROI level, enrichment scores across *ERBB2*-high tumour, tumour core, tumour boundary, adjacent stroma, ductal carcinoma in situ (DCIS)/myoepithelial, stromal/vascular and immune-enriched regions were also concordant with measured Xenium signatures; across 80 ROI-signature pairs, correlations were 0.889 for within-section reconstruction and 0.770 for cross-section transfer (Fig. 5e). These analyses move the evaluation beyond pointwise gene correlation by showing that Morpho-FM preserved boundary-level and region-level molecular organization in the tissue architecture. Full signature-level boundary-gradient profiles and ROI-level enrichment summaries are provided in Supplementary Fig. 13.

### Morpho-FM supports latent-domain discovery and candidate tissue-domain recovery across breast spatial transcriptomics platforms

We next examined whether the coupled morphology-to-transcriptome representation organized tissue before gene-specific readout. In the default Xenium S1 analysis with *k* = 15, the 1,536-dimensional pre-readout latent embeddings formed spatially contiguous domains that aligned with major tumour-rich, DCIS/myoepithelial, stromal/vascular and immune compartments in the curated tissue annotation (Fig. 6a and Supplementary Fig. 14). To assess robustness to clustering resolution and initialization, we repeated clustering for *k* = 10, 15 and 20 across five random seeds on the S1 grid-valid subset for which full-section latent features were available (Fig. 6b). Morpho-FM latent clustering consistently outperformed raw morphology clustering in coarse compartment agreement. Across *k* = 10, 15 and 20, Morpho-FM achieved normalized mutual information (NMI) values of 0.396 ± 0.004, 0.376 ± 0.006 and 0.362 ± 0.004, and adjusted Rand index (ARI) values of 0.285 ± 0.021, 0.227 ± 0.026 and 0.175 ± 0.009, respectively. Raw morphology clustering yielded substantially lower NMI values of 0.105 ± 0.007, 0.112 ± 0.009 and 0.121 ± 0.006, and ARI values of 0.070 ± 0.009, 0.057 ± 0.007 and 0.051 ± 0.003 across the same settings. Thus, the transcriptomic latent space sharpened tissue-domain structure beyond raw morphology features alone.

**Fig 6.**
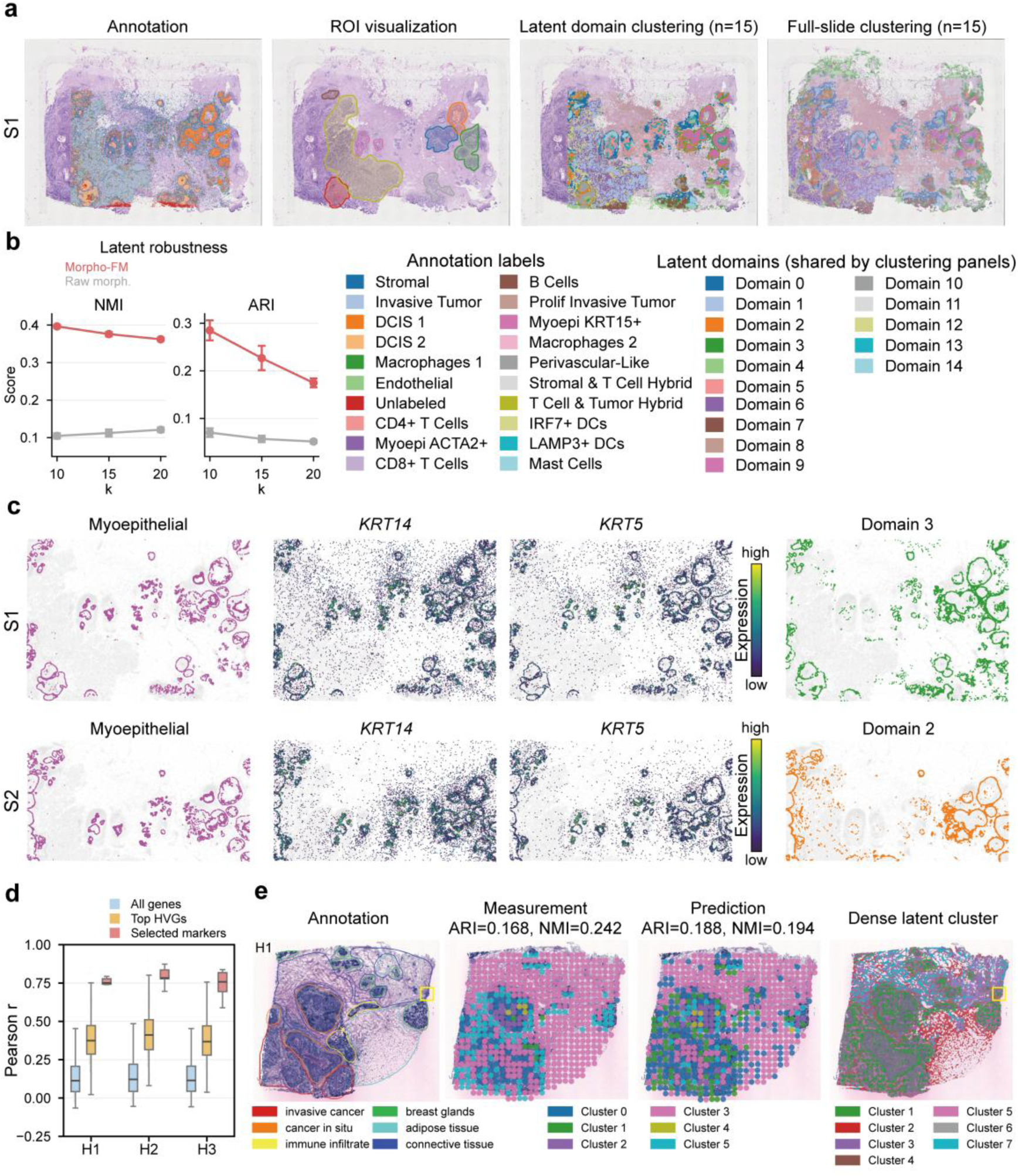
Latent-domain discovery and candidate tissue-domain recovery across breast spatial transcriptomics platforms. **a**, Curated tissue annotation, region of interest (ROI) visualization, Morpho-FM latent-domain clustering and full-section dense latent clustering in Xenium section 1 (S1). Latent domains were obtained by clustering the 1,536-dimensional pre-readout Morpho-FM latent embeddings using *k* = 15. **b**, Robustness of latent-domain clustering across *k* = 10, 15 and 20, evaluated against curated tissue annotations by normalized mutual information (NMI) and adjusted Rand index (ARI). Points and error bars indicate mean ± s.d. across five random seeds. Annotation and latent-domain colour keys are shown for the spatial maps in **a** and **c**. **c**, Spatial correspondence between myoepithelial annotations, measured *KRT14* and *KRT5* expression, and matched Morpho-FM latent domains in S1 and S2. The matched latent domains shown are domain 3 in S1 and domain 2 in S2, which spatially overlap KRT14/KRT5-enriched myoepithelial layers around DCIS structures. **d**, Leave-one-section-out prediction performance in HER2ST patient H sections. For each target section, the remaining two sections were used for training, and Pearson correlations were computed at the original HER2ST measurement locations. Boxplots summarize per-gene correlations for all retained genes, the top 1,000 highly variable genes and a selected marker panel. **e**, Label-anchored tissue-domain recovery in the held-out HER2ST H1 section. Manual annotation, measured-expression clustering, Morpho-FM predicted-expression clustering and dense latent clustering are shown for the same section. ARI and NMI quantify agreement with manual tissue labels for measured- and predicted-expression clustering at the original measurement locations. Yellow boxes indicate candidate regions not delineated in the manual annotation but identified by Morpho-FM dense latent clustering; their interpretation is supported by concordant epithelial/tumour marker expression across all three sections.

Beyond coarse compartment agreement, latent-domain analysis recovered local structures with clear histopathological relevance. In both Xenium sections, *KRT14*/*KRT5*-enriched latent domains spatially overlapped annotated myoepithelial layers, with domain 3 in S1 and domain 2 in S2 marking regions around DCIS structures (Fig. 6c). These layers formed stratified structures around DCIS regions and were depleted from invasive tumour regions. This pattern highlights a microenvironmental distinction between non-invasive and invasive breast tumour states and is consistent with established evidence that disruption of the myoepithelial cell-basement membrane interface is a defining feature of progression from in situ carcinoma to invasive breast cancer ^37,38^. Domain-level enrichment analyses further supported this interpretation, showing that DCIS/myoepithelial-enriched latent domains were concordant with curated annotations and basal/myoepithelial marker positivity across both Xenium sections. Thus, Xenium provided primary evidence that Morpho-FM can convert routine H&E morphology into a transcriptomically organized latent tissue map, not only a set of gene-wise predictions.

We then asked whether this domain-recovery workflow extended beyond Xenium to an independent breast spatial transcriptomics platform ^39^. Leave-one-section-out transfer on three consecutive HER2ST sections from patient H provided a cross-platform validation setting, with each section predicted from the remaining two; in this configuration, no target section contributed to model fitting (Supplementary Note 3). Across 13,080 retained genes, Morpho-FM achieved mean per-gene Pearson correlations of 0.140, 0.149 and 0.140 for H1, H2 and H3, respectively, with stronger prediction on the top 1,000 HVGs (mean correlations of 0.390, 0.422 and 0.383; Fig. 6d). A biologically motivated marker set spanning epithelial/tumour-associated genes (*ERBB2*, *KRT19* and *CD24*), a stromal programme marker (*MGP*) and immune-associated genes (*CD74* and *IGKC*), selected to represent major breast cancer epithelial, stromal and immune compartments described in single-cell and spatial atlases ^40,41^, showed reproducible held-out correlations across the three sections and preserved spatially organized expression patterns at the original HER2ST measurement locations (Extended Data Fig. 4). In the annotated H1 section, these marker patterns were consistent with the known tissue architecture ^39^, and similar spatial trends were observed in H2 and H3.

Finally, we evaluated whether prediction-derived representations could support tissue-domain recovery in HER2ST. In the annotated H1 section, clustering of predicted expression at the original HER2ST measurement locations showed measurable agreement with manual tissue labels (ARI/NMI, 0.188/0.194), comparable in scale to clustering of measured expression under the same setting (Fig. 6e). We further applied dense latent clustering to the whole-section latent grid, using the high-resolution representation instead of sparse measurement locations. This analysis produced coherent candidate domains aligned with the broad tissue architecture of H1 and highlighted cancer-enriched regions supported by epithelial/tumour marker fields (Fig. 6e). For the unlabelled H2 and H3 sections, dense clusters were treated as candidate domains rather than validated tissue classes. Dense latent clustering generated organized domains spatially concordant with epithelial/tumour-associated, stromal and immune marker fields in both held-out sections (Supplementary Fig. 15). Together, the Xenium and HER2ST analyses show that Morpho-FM supports candidate tissue-domain recovery across breast spatial transcriptomics platforms, while appropriately treating domain-level outputs as hypotheses for marker-guided inspection rather than independently validated tissue classes.

## Discussion

Morpho-FM addresses a central unresolved problem at the interface of computational pathology and spatial omics: whether routine H&E morphology can be decoded through a transcriptomic foundation-model prior to recover spatial molecular states when paired H&E-ST data are limited. The main contribution of this study is therefore not simply another histology-to-expression predictor, but a framework that connects two previously parallel foundation-model streams. Pathology foundation models provide scalable representations of tissue architecture, whereas single-cell transcriptomic foundation models encode structured constraints over gene-expression states. By inserting the transcriptomic foundation model at the molecular readout stage, Morpho-FM allows morphology-derived signals to be interpreted within a biologically constrained expression space rather than through an unconstrained regression head. Across prostate and kidney cancer benchmarks, external clear-cell renal cell carcinoma transfer, Xenium breast cancer reconstruction and HER2ST cross-platform analysis, this coupling improved prediction and supported spatial molecular reannotation of routine histology.

The ablation experiments sharpen this interpretation. Pretrained histology encoders are valuable because local morphology remains the conditioning signal for every prediction, but changing the upstream histology feature backbone did not produce a stable independent gain in the matched prostate controls. In contrast, pretrained CellFM initialization reproducibly improved prediction, with the larger gain observed in the single-slide setting where paired supervision was most limited. This result suggests that, under sparse H&E-ST supervision, the dominant bottleneck is not only how tissue morphology is represented, but also how the molecular output space is constrained. Treating gene expression as a high-dimensional vector learned de novo from a small number of paired sections forces the model to rediscover co-expression structure, expression distributions and cell-state dependencies from limited data. Morpho-FM instead transfers part of this structure from a pretrained single-cell transcriptomic model, helping stabilize morphology-conditioned decoding.

This design also reframes dense histology-to-expression prediction. Dense molecular maps from H&E are often visually compelling, but unmeasured positions do not have direct ground truth. We therefore linked dense full-section decoding to the original measurement support through re-aggregation, so that tissue-wide molecular reconstruction remained anchored to quantitative evaluation. In Xenium breast cancer sections, Morpho-FM recovered *ERBB2*-rich tumour compartments, preserved marker programmes spanning epithelial, stromal, vascular, proliferative and immune categories, and retained boundary- and ROI-level molecular trends after re-aggregation. The latent-domain analyses further showed that pre-readout Morpho-FM representations aligned more closely with curated tissue compartments than raw morphology features alone and recovered *KRT14*/*KRT5*-enriched myoepithelial structures around DCIS regions. These findings indicate that the transcriptomic decoder contributes not only gene-wise accuracy, but also a latent organization of tissue that is more compatible with molecular compartment structure.

At the same time, the results define the appropriate scope of the framework. Morpho-FM does not replace spatial transcriptomics, and its dense fields should not be interpreted as directly measured expression. The method still requires paired H&E and spatial transcriptomic data for training, and it inherits sequencing noise, registration uncertainty, gene-panel limitations and platform-specific biases from those measurements. Re-aggregation to the original measurement support provides an important consistency check, but it cannot validate every unmeasured grid position. Similarly, latent domains and HER2ST candidate domains should be regarded as hypotheses for marker-guided inspection, orthogonal validation or pathological review, rather than independently established tissue classes. The reduced performance in one external ccRCC section also emphasizes that cross-dataset transfer remains sensitive to tissue heterogeneity, preprocessing differences and platform effects.

These limitations point to several priorities for future development. First, broader multi- centre testing across tumour types, staining protocols, scanners and spatial transcriptomic platforms will be needed to determine the operating range of morphology-to-transcriptome decoding. Second, calibrated gene- and location-specific uncertainty estimates should accompany dense molecular maps, so that users can distinguish confident molecular reannotation from low-confidence extrapolation. Third, future models should improve cross-platform harmonization and explicitly model differences in spatial resolution, gene-panel coverage and measurement noise. Finally, the most important validation will be prospective and orthogonal: using Morpho-FM-derived molecular maps to prioritize regions for additional spatial profiling, multiplex imaging or pathological review, and then testing whether the predicted transcriptional states, spatial distributions and tumour-microenvironmental interactions are confirmed. Overall, Morpho-FM supports a shift from de novo morphology-to-transcriptome regression toward foundation-model-constrained molecular decoding from routine histology. By coupling pathology-scale morphology with a single-cell transcriptomic prior, the framework uses sparse spatial transcriptomic supervision to extend molecular information across H&E sections while preserving a clear distinction between measured data and model-inferred maps. This makes Morpho-FM relevant for molecular reannotation of pathology archives, selection of regions for orthogonal profiling and exploration of spatial gene-expression programmes in tissue regions that were not directly assayed. More broadly, the study suggests that the next step for computational pathology may not be only larger image encoders, but architectures that connect visual tissue structure to pretrained biological state spaces.

## Methods

### Study design and data preprocessing

Morpho-FM was developed to predict spatial gene expression from routine H&E whole-slide images using weak supervision from measured ST locations, and to extend the same trained model to dense full-section reconstruction. To evaluate whether this framework generalizes beyond a single tissue type, ST platform or analysis setting, we designed a study comprising multiple complementary evaluation regimes: cross-section prediction under harmonized ST benchmarks, transfer to independently processed external sections, measurement-constrained dense reconstruction in higher-resolution spatial assays and downstream analyses testing whether predicted expression or latent representations preserve biologically meaningful tissue domains. This design was intended to assess both predictive accuracy and biological utility across variations in tissue context, measurement density, platform format and train–test separation. The specific datasets, sample identifiers and experimental splits used for each evaluation regime are reported in the corresponding subsections below. All datasets analysed in this study were publicly available, and no new human tissue specimens were collected. Across all evaluation regimes, Morpho-FM requires input data to follow a unified preprocessing format. Spatial measurement coordinates were projected into H&E pixel space so that image features, measurement locations and dense prediction grids resided in a shared coordinate system. Raw expression counts were loaded as integers for the negative-binomial training objective, and model outputs were interpreted as normalized expression-rate predictions. Gene identifiers were intersected across samples and further intersected with the CellFM vocabulary to ensure that each input gene set was aligned with the pretrained representation space. The resulting gene-panel size for each dataset after this unified procedure, including 17,512 genes for the Visium-style prostate and kidney benchmarks, 306 genes for the Xenium breast cancer analysis and 13,080 genes for the HER2ST breast cancer analysis, is reported in the corresponding experimental subsections. Platform-specific adaptations required to accommodate differences in raw data formats, including AnnData count-layer naming conventions, slide-level scale-factor handling and Xenium cell-centroid coordinate conversion, are described in Supplementary Note 4.

### Whole-slide histology feature caching

Morpho-FM does not train directly from raw whole-slide pixels. Instead, each H&E whole-slide image was first converted into a cached whole-slide histology feature grid using a pretrained histology feature encoder. H&E whole-slide images were resized when necessary to limit the maximum image dimension, padded to ensure exact tiling and encoded using HIPT. The resulting feature grid was defined at one-sixteenth of the H&E pixel resolution. For each grid location, HIPT-derived local and sub-patch representations were concatenated with low-resolution red-green-blue (RGB) information to obtain a 579-dimensional morphology feature vector. Feature channels were smoothed and normalized per slide before downstream model training. All subsequent operations, including local bag construction, model training, dense decoding and re-aggregation to the original measurement support, used this cached feature grid rather than raw image patches. Feature-grid construction and normalization are detailed in Supplementary Note 5.

For comparison of histology feature backbones, a ResNet-50 feature cache was also generated in a controlled ablation. In this control, H&E images were tiled using the same cached-grid convention, ImageNet-pretrained ResNet-50 features were extracted and projected to match the downstream cache interface. All Morpho-FM components downstream of the feature cache were held fixed, allowing the analysis to isolate the effect of changing the histology feature backbone.

### Coordinate system and local disk-shaped MIL bags

Morpho-FM represents each spatial measurement location as a local disk-shaped MIL bag on the cached feature grid. Let 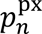 denote the H&E pixel coordinate of measurement location *n*, and let 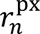 denote the corresponding measurement radius in pixel space. These quantities were mapped to the cached feature grid as

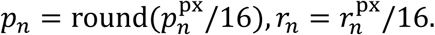

The MIL bag for location *n* was defined as the set of grid features falling within a disk centred at *p_n_*:

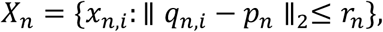

where *x_n,i_* ∈ ℝ^579^ is the cached morphology feature vector at grid position *q_n,i_* . Locations whose disk neighbourhood extended outside the valid feature grid or tissue mask were excluded before training. This formulation treats each measured spot or cell-resolved Xenium location as a weakly supervised bag: expression supervision is available only at the measurement location, whereas the model learns from a set of local histological instances within the corresponding measurement support. MIL bag construction and validity filtering are detailed in Supplementary Note 6.

### Morpho-FM architecture

Morpho-FM couples a lightweight morphology-to-transcriptome adapter to a transcriptomic decoder implemented with CellFM. Each instance feature *x_n,i_* from the local MIL bag is first projected into the CellFM latent space by a two-layer fully connected adapter with layer normalization, nonlinear activation and dropout. The adapter maps the 579-dimensional histology feature vector to the 1,536-dimensional CellFM latent space.

The projected morphology representation was used to condition transcriptomic decoding in the CellFM latent space. For each gene *g*, the decoder produced a non-negative normalized expression-rate prediction for each instance:

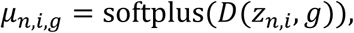

where *Z_n,i_* is the adapter output and *D*(⋅, *g*) denotes the CellFM-based gene-conditioned decoder. Instance-level rates within each bag were aggregated by unweighted mean pooling:

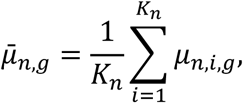

where *K_n_* is the number of grid instances in the bag. The bag-level rate μ̅_*n*,*g*_ is the primary output used for spot-level or Xenium-location-level evaluation.

During fine-tuning, most CellFM backbone parameters were frozen to preserve the pretrained transcriptomic prior. Trainable parameters were restricted to the morphology-to-transcriptome adapter, the CellFM gene embedding matrix, the decoder projection and gene-specific dispersion parameters. This parameter-efficient training design was used for all primary Morpho-FM models and was also retained in the transcriptomic initialization ablation, ensuring that pretrained and randomly initialized CellFM variants had identical trainable parameter scope. Adapter, decoder and trainable-parameter details are provided in Supplementary Note 7.

### Negative-binomial training objective

Morpho-FM was optimized using a negative-binomial count objective to account for overdispersion in ST measurements. For measurement location *n* and gene *g*, the bag-level rate μ̅_*n*,*g*_ predicted by the architecture described above was scaled by a location-specific size factor *S_n_* to obtain the count-scale mean *S_n_μ̅_n,g_* . The size factor was computed as the ratio of the total raw count at location *n* to the mean library size across the corresponding training set. The training loss was defined as

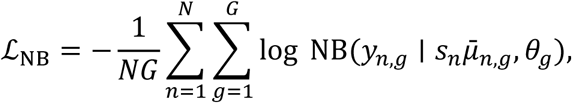

where *y_n,g_* is the raw count at location *n* for gene *g*, and *θ_g_* = exp (*ϕ_g_*) is a gene-specific dispersion parameter, with *ϕ;_g_* learned jointly with the other trainable model parameters. This formulation separates the model output, a normalized histology-conditioned expression rate, from the count-scale likelihood used for parameter estimation. For large gene panels, the loss was evaluated using shuffled gene chunks as a memory-control strategy; full implementation details are provided in Supplementary Note 8.

### Model training and checkpoint selection

All Morpho-FM models were trained using AdamW optimization with separate learning-rate groups for the transcriptomic decoder parameters and the morphology-to-transcriptome adapter and dispersion parameters. Training used automatic mixed precision, early stopping on validation negative-binomial loss and retention of the checkpoint with the lowest validation loss for all downstream inference and evaluation. The default training configuration used 100 maximum epochs, a batch size of four measurement locations, a weight decay of 10^−2^, an early-stopping patience of five epochs and a random seed of 42. Gene chunk size was set to 256 genes for large panels and was increased accordingly for smaller panels to ensure full coverage within fewer forward passes. Complete hyperparameters, learning-rate schedules and implementation details are provided in Supplementary Note 8.

The inference workflow accepted an explicit checkpoint and target section as inputs, enabling both within-section dense reconstruction and cross-section transfer without modifying the training procedure or the dense-decoding pipeline.

### Dense full-section reconstruction and re-aggregation

After spot-supervised or location-supervised training, the same adapter-conditioned transcriptomic decoder was applied to every valid grid position under the tissue mask to generate a dense expression-rate field,

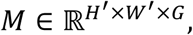

where *H*^′^ and *W*^′^ denote the cached feature-grid dimensions and *G* is the number of genes being decoded. Dense inference did not apply spot-specific library-size scaling; therefore, *M* was interpreted as a normalized spatial expression-rate field rather than an absolute count matrix. Grid positions outside the tissue mask or outside the finite feature region were excluded from downstream analyses.

To assess whether dense predictions remained consistent with measured expression at the original measurement support, dense predictions were re-aggregated to the measurement support using the same disk kernel used for MIL bag construction. For location *n* and gene *g*, the re-aggregated dense prediction was computed as the mean of *M_u,v,g_* over the grid positions (*u, v*) within the disk centred at *p_n_*. The Pearson correlation between the re-aggregated dense prediction and measured expression at the original measurement support was used as an internal consistency measure for dense reconstruction. This re-aggregation analysis does not constitute direct validation at unmeasured grid locations; rather, it tests whether dense full-section predictions remain coupled to the measured expression signal when projected back to the measurement scale. Dense decoding and re-aggregation are detailed in Supplementary Note 9.

### Prostate benchmark protocol

The prostate benchmark compared Morpho-FM with five representative histology-to-expression methods: HisToGene, iStar, mclSTExp, sCellST and HiST. Four HEST prostate cancer sections, INT25, INT26, INT27 and INT28, were used. Each section comprised an H&E whole-slide image, an ST count matrix, spatial measurement coordinates and assay scale-factor metadata. All methods were evaluated on the same sections, the same H&E-aligned spatial coordinates and the same 17,512-gene panel. Prediction outputs from all methods were converted to a common spot-ordered and gene-ordered matrix before evaluation. Method-specific input conversion, validation handling and output alignment were implemented as needed, but the core architectures and loss functions of the baseline methods were not modified.

Two prostate benchmark protocols were used. In the rotating single-slide protocol, each of the four prostate sections served once as the sole training slide, and the remaining three sections were held out for independent testing. This yielded 12 directed train–test comparisons. Validation in the single-slide setting was constructed within the training slide using a spatially blocked partition rather than a random spot split, reducing leakage from local spatial autocorrelation. In the rotating multi-slide protocol, two prostate sections were used for training, one section for validation and one section for held-out testing. The held-out test section was rotated across the four prostate sections, yielding four folds. The validation slide or blocked validation subset was used only for checkpoint selection and early stopping.

### Kidney benchmark and external ccRCC transfer

The kidney HEST benchmark used the same harmonized evaluation framework as the prostate benchmark. Eight HEST kidney sections, INT13, INT14, INT15, INT17, INT18, INT19, INT21 and INT24, were evaluated under a rotating single-slide design, yielding 56 directed train–test comparisons. All methods were aligned to the shared kidney gene panel after CellFM vocabulary mapping, and performance was evaluated using the same per-gene correlation metrics as in the prostate benchmark.

For external transfer, a fixed multi-slide kidney setting was used to select a kidney-trained Morpho-FM checkpoint. Six HEST kidney sections were used for training, INT24 was used for validation and INT21 was used as an internal held-out test section. The resulting checkpoint was then applied without further fitting, target-section-specific tuning or hyperparameter adjustment to two external ccRCC Visium sections. External sections were converted to the same Morpho-FM input interface, including count-matrix processing, gene harmonization, coordinate handling and HIPT feature-cache generation. Transfer performance was evaluated on the shared gene panel using per-gene Pearson correlation, global expression correlation, mean spot-level expression-profile correlation and marker-level spatial inspection.

### Transcriptomic initialization and histology feature-backbone ablations

To isolate the contribution of pretrained transcriptomic initialization, we performed a controlled 2 × 2 ablation crossing two training regimes with two initialization conditions. The training regimes were a multi-slide prostate setting and a single-slide prostate setting. The initialization conditions were pretrained CellFM weights and randomly initialized CellFM weights. All four branches used the same data partitions, gene panel, MIL bag geometry, negative-binomial objective, trainable parameter scope, training budget, early-stopping procedure and inference pipeline. Performance gain from pretrained transcriptomic initialization was defined as the difference in the chosen evaluation metric between the pretrained and randomly initialized CellFM branches under the same training regime.

To evaluate the influence of the histology feature backbone, a separate encoder-control ablation replaced the HIPT feature cache with an ImageNet-pretrained ResNet50 feature cache while keeping the pretrained CellFM branch and all downstream Morpho-FM components fixed. This control was evaluated under matched prostate single-slide and multi-slide settings. This design allowed the ablation analysis to distinguish the contribution of pretrained transcriptomic initialization from the effect of replacing the cached histology feature representation.

### Xenium breast reconstruction analysis

The Xenium analysis used two consecutive 10x Genomics Xenium FFPE breast cancer sections, S1/XEN1 and S2/XEN2. Xenium cell centroids were mapped into H&E coordinates using the vendor-provided morphology-to-H&E alignment information, and an effective location diameter was estimated from cell area so that the same disk-based geometric framework could be reused. The effective measurement radius was used to define local histological neighbourhoods and re-aggregation support. After intersection with the CellFM vocabulary and across-section harmonization, 306 genes were retained for downstream analysis.

Four reconstruction and transfer settings were evaluated: S1-to-S1, S1-to-S2, S2-to-S2 and S2-to-S1. The within-section settings were interpreted as measurement-constrained dense reconstruction analyses because the same section provided the measured locations used to constrain the model; these settings were not interpreted as cross-section generalization tests. The reciprocal cross-section settings tested whether the learned morphology-to-expression mapping transferred between consecutive sections. For each setting, predictions were evaluated at observed Xenium measurement locations and, after dense full-section decoding, by re-aggregation to the original Xenium measurement support.

To avoid relying on a single marker, we evaluated both the full 306-gene panel and a fixed 29-gene breast marker panel spanning tumour/luminal, epithelial, proliferative, stromal, vascular and immune programmes. *ERBB2* was used as a well-characterized visualization marker, but section-level conclusions were based on the full-gene and balanced marker-panel analyses. Boundary-gradient analyses used curated tissue-compartment annotations to define tumour-adjacent spatial bins by signed distance to the tumour edge. ROI-level analyses compared measured and predicted signature enrichment across curated regions, including *ERBB2*-high tumour, tumour core, tumour boundary, adjacent stroma, DCIS/myoepithelial, stromal/vascular and immune-enriched areas. The Xenium reconstruction settings, marker panel and region-level analyses are detailed in Supplementary Note 10.

### Xenium latent-domain analysis

To determine whether Morpho-FM captured tissue organization before gene-specific readout, we extracted the 1,536-dimensional pre-readout Morpho-FM latent representation, corresponding to the CellFM latent space, before the final gene decoder. Latent vectors were computed at valid Xenium measurement locations and reduced by principal component analysis. MiniBatch K-means clustering was then applied to the reduced latent space, with *k* = 15 used as the default clustering resolution. The fitted dimensionality-reduction and clustering models were projected to all valid grid positions in the tissue mask to generate dense full-section latent-domain maps whose cluster identities were aligned with the measurement-location latent representation.

Robustness to clustering resolution and initialization was evaluated by repeating latent clustering for *k* = 10, *k* = 15 and *k* = 20 across five random seeds. Agreement with curated tissue-compartment annotations was quantified using NMI and ARI. Raw morphology clustering was used as a baseline to determine whether Morpho-FM latent representations provided more transcriptomically aligned tissue-domain structure than morphology features alone. Domain-level biological interpretation was performed post hoc by cross-tabulating latent domains with curated annotations and by assessing enrichment of measured basal/myoepithelial markers, including *KRT14* and *KRT5*. These annotations and marker enrichments were not used during model training, dense inference or unsupervised clustering.

### HER2ST breast-platform support analysis

The HER2ST patient H analysis was designed as a supporting breast-platform testbed rather than as a co-primary validation cohort. Sections H1, H2 and H3 were converted into the same Morpho-FM input format used in the Visium and Xenium analyses, including H&E image input, spatial coordinates, count matrices and slide metadata. After gene harmonization across H1–H3 and CellFM vocabulary mapping, 13,080 genes were retained.

HER2ST prediction was evaluated using a leave-one-section-out transfer design. For each target section, the remaining two sections were used for model fitting and checkpoint selection, and the target section was excluded from training and validation. Checkpoint selection used validation data drawn only from the training sections. Thus, H1 was predicted from models trained on H2 and H3, H2 from H1 and H3, and H3 from H1 and H2. Prediction performance was summarized at the original HER2ST measurement locations across all retained genes, across the top 1,000 HVGs and across a selected epithelial, stromal and immune marker panel.

To test whether predicted expression retained sufficient structure for downstream domain recovery, measured and predicted expression profiles were clustered at the original HER2ST measurement locations. Section H1, the only section with available tissue labels in the released metadata, was used for label-agreement evaluation with ARI and NMI. Whole-section dense latent clustering was then applied to the held-out target sections to generate higher-resolution candidate domains from the Morpho-FM latent representation. Sections H2 and H3 were treated as unlabelled exploratory sections; their dense latent clusters and marker maps were used to generate candidate spatial domains for marker-based inspection rather than validated tissue classes. Whole-section dense decoding was performed for selected variable and marker genes to visualize epithelial, stromal and immune-associated spatial patterns.

### Evaluation metrics

The primary prediction metric was per-gene Pearson correlation between predicted and measured expression across spatial locations on the held-out test section or target section. Genes with zero variance or undefined correlation were excluded from correlation summaries. For each experiment, per-gene correlations were summarized by mean and median values across the retained gene panel. For external kidney transfer, we also reported global expression correlation across the full location-by-gene matrix and mean spot-level expression-profile correlation across locations.

HVG analyses were performed post hoc and were not used for training, checkpoint selection or hyperparameter tuning. For HVG subset evaluation, HVGs were identified from the measured expression matrix in the relevant test section. Prediction quality on the top 1,000 measured HVGs was summarized by mean per-gene Pearson correlation. For HVG recovery analysis, top- *K* predicted variable genes were compared with measured HVGs across *K* cutoffs, and Recall@*K* was reported together with the mean Pearson correlation of the predicted top-*K* gene set.

For dense full-section reconstruction, evaluation was restricted to the original measurement support after re-aggregation because unmeasured grid locations do not have direct ground truth. Dense-to-measurement re-aggregation Pearson correlation was therefore interpreted as an internal consistency metric rather than as direct validation of unmeasured positions. For clustering analyses, agreement between inferred domains and curated annotations was quantified using NMI and ARI. Marker-level and ROI-level analyses were used as biological consistency checks rather than as independent supervised training signals. Metric definitions and finite-value aggregation rules are detailed in Supplementary Note 11.

## Data availability

All spatial transcriptomics datasets analysed in this study are publicly available. The primary source for prostate and kidney sections was the HEST collection hosted on Hugging Face (MahmoodLab/hest), which also provides cell segmentation outputs: https://huggingface.co/datasets/MahmoodLab/hest. The Xenium breast cancer data were obtained from the 10x Genomics Xenium in situ preview dataset: https://www.10xgenomics.com/products/xenium-in-situ/preview-dataset-human-breast. The HER2ST patient H breast cancer sections (H1–H3) were downloaded from Zenodo: https://zenodo.org/records/4751624. The two external ccRCC Visium sections used for transfer evaluation were obtained from Davidson *et al.* (GEO accession GSE210038).

## Code availability

Code for Morpho-FM and the harmonized benchmark evaluation workflows is available at https://github.com/StickTaTa/Morpho-FM. Detailed implementation settings, feature-cache construction, model hyperparameters, baseline adaptations, split definitions, ablation protocols and evaluation scripts are described in Supplementary Notes 1-12. Reproducibility and auditability are summarized in Supplementary Note 12.

## Acknowledgements

We acknowledge computing support from the National Supercomputer Center in Guangzhou (Tianhe-2A) and from the High-performance Computing Public Platform (Shenzhen Campus) of Sun Yat-sen University.

## Author contributions

J.H. developed the method, designed and performed the experiments, analysed the data, prepared the figures and wrote the manuscript. X.F. and L.Q. assisted with data processing, benchmark evaluation and interpretation of the results. L.Z. conceived and supervised the project, provided scientific guidance and revised the manuscript. All authors reviewed and approved the final manuscript.

---

**Extended Data Fig 1.**
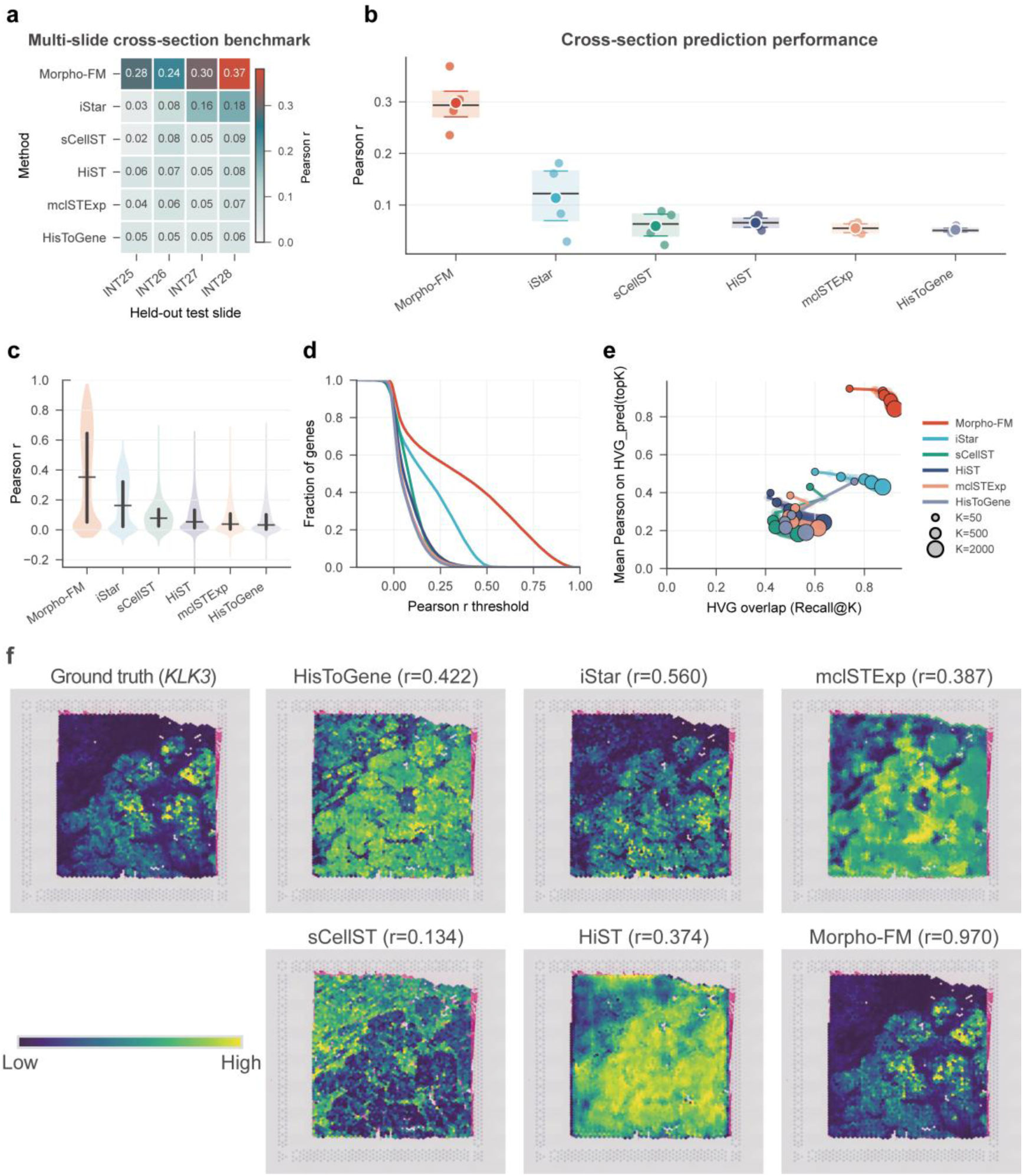
Multi-slide cross-section benchmarking of Morpho-FM on prostate spatial transcriptomics data. a, Rotating multi-slide train–validation–test design across four prostate cancer sections. In each fold, two sections were used for training, a third section was used for validation and the remaining section was held out for testing. Heatmap values indicate mean per-gene Pearson correlation for each method on each held-out test section. **b,** Mean per-gene Pearson correlation across the four held-out folds for Morpho-FM and competing methods. Points denote individual held-out folds, and boxplots summarize the distribution across folds. **c,** Per-gene Pearson correlation distributions for the representative INT26+INT27-to-INT28 fold, with INT25 used for validation. **d,** Fraction of genes with Pearson correlation above each threshold in the representative INT26+INT27-to-INT28 fold. **e,** HVG recovery and prediction quality across top-*K* predicted gene sets in the representative fold. Recall@*K* measures overlap with highly variable genes, mean Pearson correlation is computed for the predicted top-*K* gene set, and point size denotes *K* = 50, 200, 500, 1,000 or 2,000. **f,** Spatial visualization of *KLK3* expression in the representative INT26+INT27-to-INT28 fold. Ground truth and method predictions are shown after aggregation to the original measurement locations; values in parentheses denote Pearson correlation coefficients between predicted and measured *KLK3* expression.

**Extended Data Fig 2.**
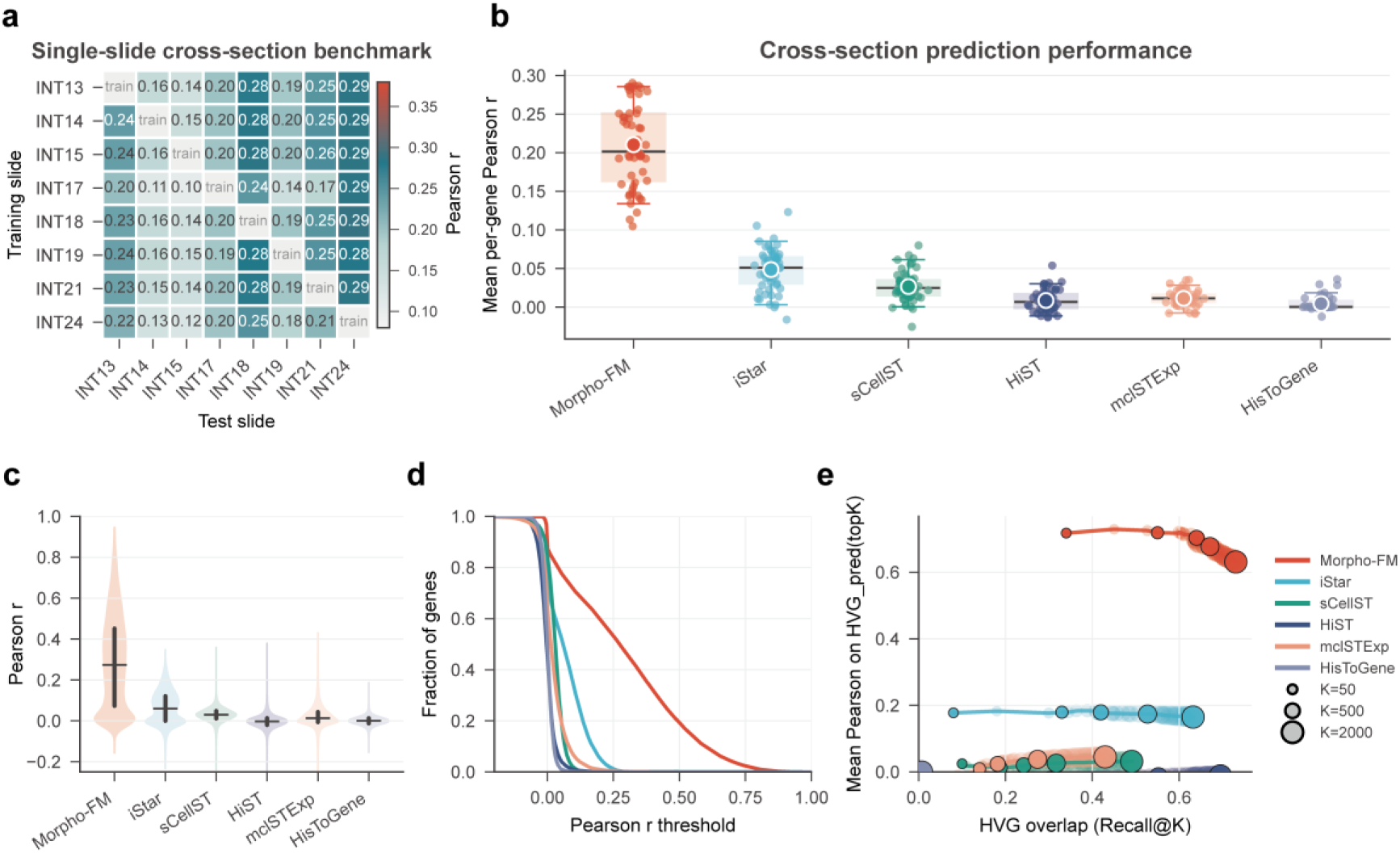
Single-slide cross-section benchmarking of Morpho-FM on kidney cancer spatial transcriptomics data. a, Rotating single-slide train–test design across eight HEST kidney cancer sections. In each train–test direction, one section was used for training and a different section was held out for testing, yielding 56 train–test directions. Heatmap values indicate mean per-gene Pearson correlation for each training and test section pair. **b,** Mean per-gene Pearson correlation across the 56 kidney cancer train–test directions for Morpho-FM and competing methods. Points denote individual train–test directions, and boxplots summarize the distribution across directions. **c,** Per-gene Pearson correlation distributions across the kidney cancer single-slide benchmark for Morpho-FM and competing methods. **d,** Fraction of genes with Pearson correlation above each threshold in the kidney cancer single-slide benchmark. **e,** HVG recovery and prediction quality across top-*K* predicted gene sets in the kidney cancer single-slide benchmark. Recall@*K* measures overlap with highly variable genes, mean Pearson correlation is computed for the predicted top-*K* gene set, and point size denotes *K* = 50, 200 or 1,000.

**Extended Data Fig 3.**
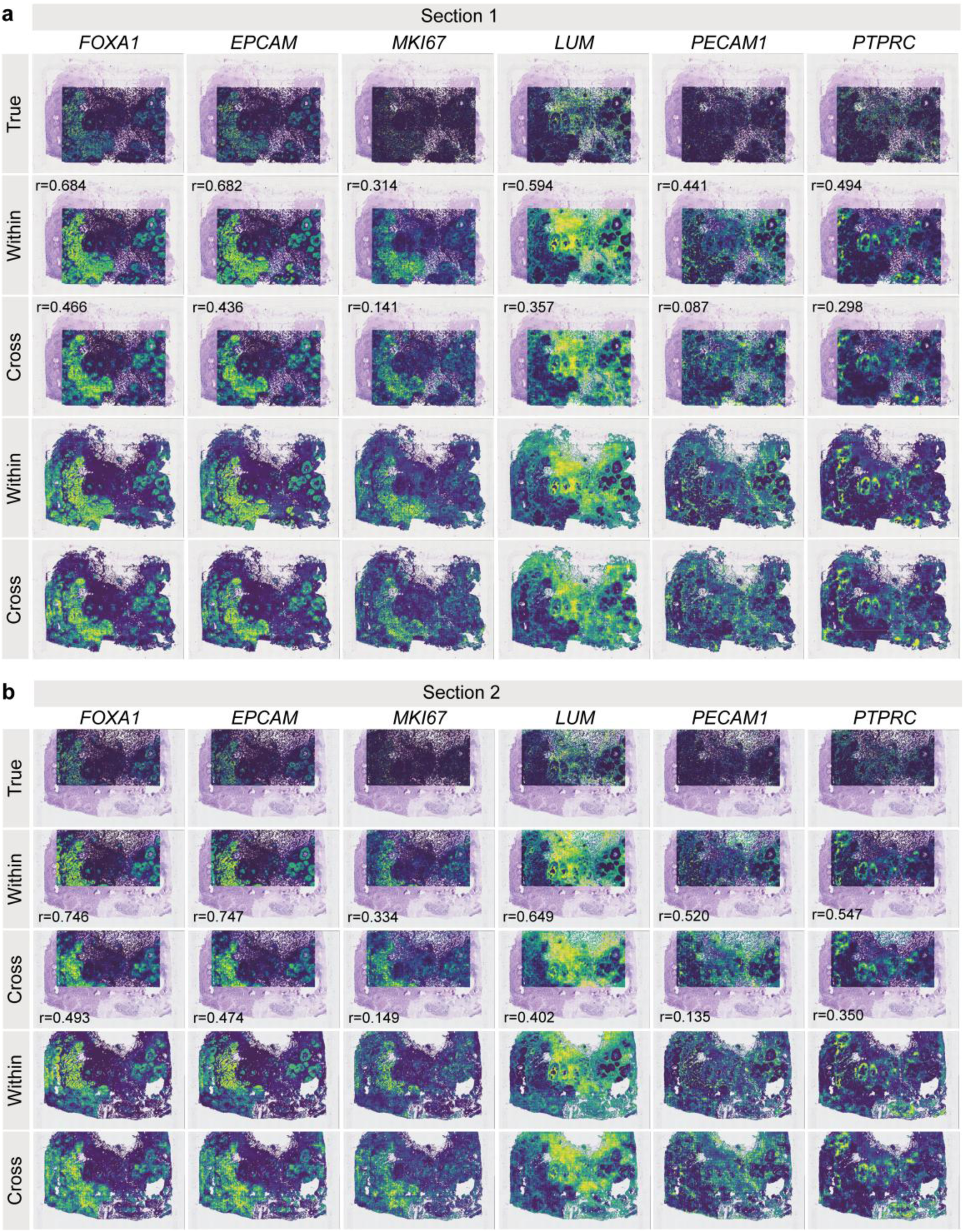
Representative marker-class Xenium prediction and dense reconstruction maps. a,b, Representative genes spanning the six marker classes in Xenium section 1 (**a**) and section 2 (**b**): *FOXA1* for tumour/luminal, *EPCAM* for epithelial, *MKI67* for proliferation, *LUM* for stromal, *PECAM1* for vascular and *PTPRC* for immune programmes. For each section, rows show measured Xenium expression at observed measurement locations, within-section prediction at the same locations, reciprocal cross-section prediction at the same locations, within-section dense full-section reconstruction and reciprocal cross-section dense full-section reconstruction. Pearson correlation coefficients are shown for the observed-location prediction rows. These examples provide spatial marker-level context for the 29-gene panel summaries and illustrate that Morpho-FM preserves major tissue-scale expression patterns across multiple biological programmes, with stronger within-section reconstruction and attenuated but visible reciprocal transfer.

**Extended Data Fig 4.**
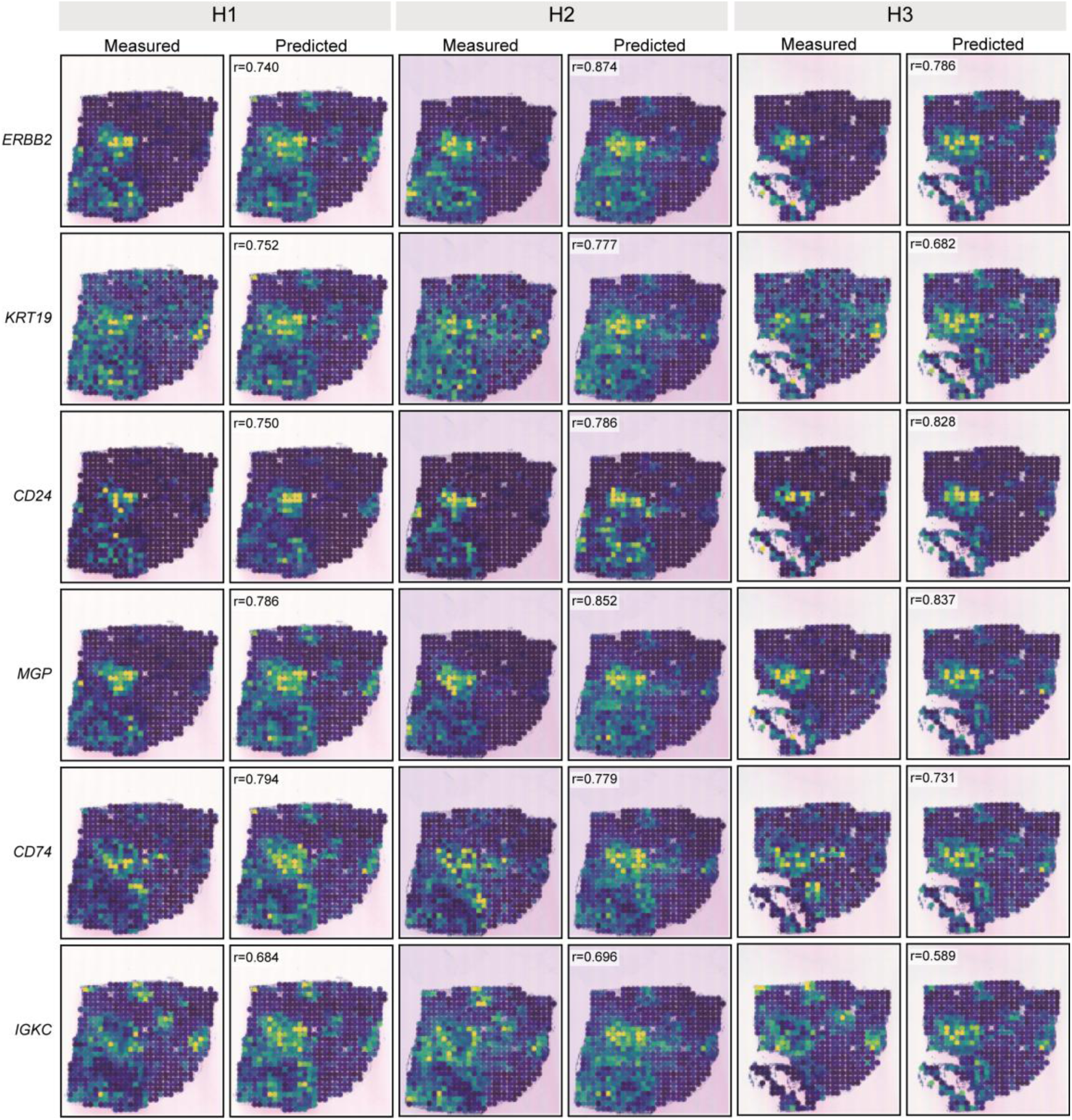
Spot-level marker prediction in held-out HER2ST patient H sections. Measured and Morpho-FM-predicted expression maps are shown for six representative marker genes across three HER2ST patient H sections. For each target section, leave-one-section-out transfer was used: H1 was predicted from H2 and H3, H2 from H1 and H3, and H3 from H1 and H2, so the target section was not used for model fitting. Rows show epithelial/tumour-associated markers (*ERBB2*, *KRT19* and *CD24*), a stromal-associated marker (*MGP*) and immune-associated markers (*CD74* and *IGKC*). For each section and gene, measured expression at the original HER2ST measurement locations is shown next to the corresponding predicted expression. Pearson correlation coefficients between measured and predicted spot-level expression are indicated in the predicted panels. Colour intensity denotes relative expression within each map.

