## Supplementary Note for "Morpho-FM: spatial molecular reconstruction from routine H&E histology using transcriptomic foundation-model priors"

Extended Computational Methods for Morpho-FM

**3 Supplementary Note 3: HER2ST Breast-Platform Support Analysis** **5**

**4 Supplementary Note 4: Notation and Data Representation** **5**

**5 Supplementary Note 5: Whole-Slide Histology Feature Grid** **6**

**8 Supplementary Note 8: Negative-Binomial Training Objective 9**

**9 Supplementary Note 9: Dense Decoding and Re-Aggregation 9**

**10 Supplementary Note 10: Xenium Breast Reconstruction Analysis 10**

**12 Supplementary Note 12: Reproducibility and Auditability 12**

### Purpose of This Note

This Supplementary Note provides an expanded methodological description of Morpho-FM. The main manuscript presents the biological motivation, principal experiments and interpretation. Here, we give the computational details needed to understand how haematoxylin and eosin (H&E) histology features, spatial measurement coordinates, CellFM gene embeddings, negative-binomial training and dense re-aggregation are connected in one framework. The note also defines the harmonized benchmark protocol used to compare Morpho-FM with existing histology-to-expression methods [1, 2, 3, 4, 5, 7].

No figures are included in this note. Tables are used only where they compactly define datasets, splits, marker panels or hyperparameters.

### 1 Supplementary Note 1: Benchmark Design

#### 1.1 Fairness criteria

All benchmark comparisons were designed around four controls. First, all methods were evaluated on matched train/validation/test slide definitions whenever the method interface allowed it. Second, all predictions were aligned to the same retained gene set for the relevant cohort. Third, method-specific output formats were converted to a common location-by-gene matrix before metric computation. Fourth, all metrics were recomputed using the same evaluation functions, rather than relying on method-specific reporting conventions.

#### 1.2 Prostate splits

The prostate benchmark used four HEST sections, INT25–INT28. The single-slide protocol used one training section and evaluated all remaining sections, yielding 12 directed train-test comparisons. The multi-slide protocol used two training sections, one validation section and one held-out test section, yielding four folds.

Table 1: Prostate benchmark partitions.

| Protocol | Train | Validation | Test |
|---|---|---|---|
| Single-slide | one of INT25–INT28 | blocked split within train slide | remaining three slides |
| Multi-slide | two slides | one independent slide | one held-out slide |

Across the 12 prostate single-slide directions, Morpho-FM achieved a mean per-gene Pearson correlation of 0.286. Across the four multi-slide folds, Morpho-FM achieved a mean per-gene Pearson correlation of 0.298.

##### 1.3 Kidney benchmark and external transfer

The kidney benchmark used eight HEST sections: INT13, INT14, INT15, INT17, INT18, INT19, INT21 and INT24. The rotating single-slide design used each section once as the sole training section and evaluated the remaining seven sections, yielding 56 directed comparisons. Morpho-FM achieved a mean per-gene Pearson correlation of 0.210 across these comparisons.

For external ccRCC transfer, a fixed kidney-trained checkpoint was selected using a multi-slide split with INT21 as the internal held-out test section. The internal INT21 mean per-gene Pearson correlation was 0.255. The same checkpoint was then applied, without target-section fine-tuning, to two external ccRCC Visium sections. The corresponding mean per-gene Pearson correlations were 0.294 and 0.138 for external sections 1 and 2, respectively.

##### 1.4 Baseline methods

Five finalized baseline methods were included: HisToGene, iStar, mclSTExp, sCellST and HiST [1, 2, 3, 4, 5]. Baseline adaptations were restricted to data interface conversion, split handling, checkpoint selection and output alignment. Core model architectures and objectives were not altered. An exploratory THItoGene adaptation was attempted during benchmark development [6], but it was not included in final quantitative comparisons because reliable gene-aligned outputs were not obtained under the harmonized protocol.

#### 2 Supplementary Note 2: CellFM Prior and Histology-Encoder Ablations

##### 2.1 CellFM prior ablation

The CellFM prior was evaluated with a $2 \times 2$ ablation crossing training regime and initialization:

$$\Delta^{(a)}_{\text{prior}} = M^{(a)}_{\text{pretrained}} - M^{(a)}_{\text{random}},$$

where $a$ denotes the training regime and $M$ is the evaluation metric, such as mean per-gene Pearson correlation. In the multi-slide setting, pretrained CellFM initialization achieved 0.360 compared with 0.299 for random initialization. In the single-slide setting, the corresponding values were 0.346 and 0.265. Because trainable parameter scope, data partitions, MIL geometry, loss function and optimization settings were held fixed, these deltas estimate the contribution of the pretrained transcriptomic initialization under matched conditions.

##### 2.2 Expression-stratified prior analysis

To determine whether the CellFM prior acted uniformly across genes, genes were stratified by measured expression level. Let $\bar{y}_g = N^{-1} \sum_n y_{n,g}$ be the mean expression of gene $g$ in the relevant training setting. Genes were binned into low-, mid- and high-expression strata according to quantiles of $\log(1 + \bar{y}_g)$. For each bin $b$, the prior gain was computed as

$$\Delta_b = \frac{1}{|\mathcal{G}_b|} \sum_{g \in \mathcal{G}_b} \left( r_g^{\text{pretrained}} - r_g^{\text{random}} \right).$$

This analysis showed that gains were concentrated in genes with detectable measured expression structure, supporting the interpretation that CellFM provides a transcriptomic co-expression constraint rather than a uniform numerical offset.

### 3 Supplementary Note 3: HER2ST Breast-Platform Support Analysis

HER2ST patient H sections H1, H2 and H3 were used as an independent breast spatial transcriptomics platform. After gene harmonization and CellFM vocabulary mapping, 13,080 genes were retained. Leave-one-section- out transfer was used: for each target section, the remaining two sections were used for model fitting and checkpoint selection, and the target section was excluded from training and validation. Performance was summarized across all retained genes and the top 1,000 highly variable genes (HVGs).

Table 2: HER2ST leave-one-section-out prediction summary.

| Target | Training sections | All-gene mean $r$ | HVG mean $r$ |
|---|---|---|---|
| H1 | H2 + H3 | 0.140 | 0.390 |
| H2 | H1 + H3 | 0.149 | 0.422 |
| H3 | H1 + H2 | 0.140 | 0.383 |

The HER2ST marker inspection panel contained epithelial/tumour-associated genes (ERBB2, KRT19 and CD24), a stromal marker (MGP), and immune- associated genes (CD74 and IGKC). In H1, the only patient H section with available manual labels in the processed metadata, measured-expression and predicted-expression clustering were compared with manual labels at the original measurement locations. Predicted-expression clustering reached ARI/NMI 0.188/0.194. For H2 and H3, dense latent clusters were treated as candidate domains for marker-based inspection rather than validated tissue classes.

### 4 Supplementary Note 4: Notation and Data Representation

For a tissue section $s$, let $I_s$ denote the H&E whole-slide image, $Y_s \in \mathbb{N}^{N_s \times G_s}$ the raw spatial transcriptomics (ST) count matrix, and $C_s = \{c_{s,n}\}_{n=1}^{N_s}$ the spatial measurement coordinates in the assay coordinate system. Each measurement location $n$ has a count vector $y_{s,n}$ and an associated H&E pixel coordinate $p_{s,n}^{px}$. The mapping from assay coordinates to H&E pixels is written as

$$p_{s,n}^{px} = T_s(c_{s,n}),$$

where $T_s$ is the section-specific coordinate transform. For Visium-style HEST-1k (HEST) sections [8], $T_s$ is determined by the slide scale-factor metadata. For Xenium, $T_s$ combines conversion from morphology-image units to pixels with the vendor-provided morphology-to-H&E alignment transform. For HER2ST [9] and external clear-cell renal cell carcinoma (ccRCC) sections [10], the processed coordinates were converted into the same H&E pixel coordinate convention before feature extraction.

The retained gene set for an experiment is

$$\mathcal{G} = \left( \bigcap_{s \in \mathcal{S}} \mathcal{G}_s \right) \cap \mathcal{V}_{\text{CellFM}},$$

where $\mathcal{S}$ is the set of sections included in the experiment, $\mathcal{G}_s$ is the detected gene set in section $s$, and $\mathcal{V}_{\text{CellFM}}$ is the CellFM vocabulary [15]. Each retained gene $g \in \mathcal{G}$ is mapped to its CellFM vocabulary index $m(g)$. This explicit intersection and vocabulary mapping prevents gene-order differences across sections or methods from contributing to benchmark results.

Table 3: Datasets and retained gene spaces used in Morpho-FM analyses.

| Analysis | Platform | Sections | Retained genes |
|---|---|---|---|
| Prostate benchmark | HEST ST | 4 | 17,512 |
| Kidney benchmark | HEST ST | 8 | 17,512 |
| External ccRCC transfer | 10x Visium | 2 | kidney-shared |
| Xenium breast | Xenium | 2 | 306 |
| HER2ST breast | HER2ST | 3 | 13,080 |

The corresponding section identifiers were INT25–INT28 for prostate, INT13, INT14, INT15, INT17, INT18, INT19, INT21 and INT24 for kidney, S1/XEN1 and S2/XEN2 for Xenium, and H1–H3 for HER2ST.

### 5 Supplementary Note 5: Whole-Slide Histology Feature Grid

#### 5.1 Feature extraction

Each H&E whole-slide image is converted into an offline feature grid before Morpho-FM training. Let $F_s \in \mathbb{R}^{H'_s \times W'_s \times d}$ denote the grid for section $s$, where the grid stride is 16 H&E pixels and $d = 579$. The 579 channels are a concatenation of patch-level Hierarchical Image Pyramid Transformer (HIPT) features, sub-patch HIPT features and downsampled red-green-blue (RGB) features [11]:

$$F_s(u, v) = \left[ F_s^{\text{cls}}(u, v), F_s^{\text{sub}}(u, v), F_s^{\text{rgb}}(u, v) \right],$$

with dimensions 384, 192 and 3, respectively. The feature tensor is defined on the H&E pixel coordinate system, so every spatial measurement location can be mapped to the same grid.

#### 5.2 Feature normalization

Feature channels are normalized independently within each section:

$$\hat{F}_{s,u,v,c} = \frac{F_{s,u,v,c} - \mu_{s,c}}{\sigma_{s,c} + \epsilon},$$

where

$$\mu_{s,c} = \frac{1}{|\Omega_s|} \sum_{(u,v)\in\Omega_s} F_{s,u,v,c}, \qquad \sigma^2_{s,c} = \frac{1}{|\Omega_s|} \sum_{(u,v)\in\Omega_s} (F_{s,u,v,c} - \mu_{s,c})^2.$$

Here $\Omega_s$ is the valid feature-grid domain and $\epsilon = 10^{-12}$. This normalization removes slide-level scale differences in the cached histology representation while preserving the relative spatial structure within a section.

##### 5.3 Histology-encoder control

The primary model uses the HIPT-derived feature grid. A controlled histology-encoder ablation replaced this feature grid with an ImageNet-pretrained ResNet-50 feature grid [12] while keeping all downstream components fixed: the same spatial coordinates, multiple-instance learning (MIL) bag geometry, CellFM decoder, training objective, train/validation/test split definitions and evaluation metrics. This control isolates the effect of the feature substrate from the effect of the CellFM transcriptomic prior.

#### 6 Supplementary Note 6: Disk-Shaped Multiple-Instance Learning Bag Construction

Morpho-FM uses spatial transcriptomics supervision only at measured locations. Each measured location is therefore represented as a disk-shaped multiple-instance learning (MIL) bag on the feature grid. The H&E pixel coordinate and effective measurement radius are first mapped to feature-grid coordinates:

$$\tilde{p}_{s,n} = \mathrm{round}(p^{px}_{s,n}/16), \qquad \tilde{r}_s = r^{px}_s/16.$$

The disk kernel is

$$\mathcal{D}(\tilde{r}_s) = \{(i,j) \in \mathbb{Z}^2 : i^2 + j^2 \le \tilde{r}_s^2\}.$$

The MIL bag for location $n$ is then

$$\mathcal{B}_{s,n} = \left\{ \hat{F}_s(\tilde{p}_{s,n,1} + i, \tilde{p}_{s,n,2} + j) : (i,j) \in \mathcal{D}(\tilde{r}_s) \right\}.$$

If $K_{s,n} = |\mathcal{B}_{s,n}|$, the bag can be written as $\mathcal{B}_{s,n} = \{x_{s,n,i}\}_{i=1}^{K_{s,n}}$, where $x_{s,n,i} \in \mathbb{R}^{579}$.

A measurement location is retained only if its full disk neighbourhood falls inside the valid feature grid and all required feature vectors are finite. This same validity rule is used during training and during re-aggregation of dense predictions, so the assay-support comparison is geometrically matched to the training construction.

### 7 Supplementary Note 7: Morpho-FM Architecture

#### 7.1 Visual adapter

Each instance feature $x_{s,n,i}$ is projected to the CellFM embedding dimension through a two-layer adapter:

$$z_{s,n,i} = W_2 \operatorname{Dropout}\left[\operatorname{ReLU}\left(\operatorname{LayerNorm}(W_1 x_{s,n,i} + b_1)\right)\right] + b_2,$$

where $z_{s,n,i} \in \mathbb{R}^{1536}$. This adapter is the only module that directly maps histology features into the transcriptomic foundation model embedding space.

#### 7.2 CellFM conditioning

CellFM was used as the pretrained transcriptomic prior because it was trained on a large human single-cell transcriptomic corpus and follows the broader single-cell foundation-model direction established by models such as scGPT and Geneformer [13, 14, 15].

For a gene subset $\mathcal{C} \subseteq \mathcal{G}$, let

$$E_{\mathcal{C}} = \{e_g = E[m(g)] : g \in \mathcal{C}\}$$

be the corresponding CellFM gene embeddings. The adapter output $z_{s,n,i}$ is used as a morphology-conditioned prompt token, and the CellFM retention backbone receives the sequence

$$U_{s,n,i,\mathcal{C}} = [z_{s,n,i}, e_{g_1}, e_{g_2}, \ldots, e_{g_{|\mathcal{C}|}}].$$

After CellFM propagation, the prompt output is denoted $h_{s,n,i} \in \mathbb{R}^{1536}$, and the contextualized gene embedding for gene $g$ is denoted $\tilde{e}_g$.

#### 7.3 Gene-conditioned readout

The CellFM cell-wise decoder produces an instance-level normalized expression rate for each gene:

$$\rho_{s,n,i,g} = \operatorname{softplus}\left(\langle \sigma(W_{\text{dec}} \tilde{e}_g), h_{s,n,i} \rangle\right),$$

where $W_{\text{dec}}$ is the decoder projection and $\sigma$ is the sigmoid function. The inner product makes the prediction depend on both the tissue morphology-conditioned prompt representation and the CellFM gene embedding.

#### 7.4 Bag-level aggregation

Instance-level predictions within a disk-shaped bag are aggregated by unweighted mean pooling:

$$\bar{\rho}_{s,n,g} = \frac{1}{K_{s,n}} \sum_{i=1}^{K_{s,n}} \rho_{s,n,i,g}.$$

The bag-level rate $\bar{\rho}_{s,n,g}$ is the normalized expression-rate prediction evaluated at the original measurement location.

##### 7.5 Trainable parameter scope

During fine-tuning, most CellFM backbone parameters are frozen. The trainable parameters are the visual adapter, the CellFM gene embedding matrix, the decoder projection and the per-gene dispersion parameters. The same trainable parameter scope is used in the random-initialization CellFM ablation, so the comparison tests the pretrained transcriptomic prior rather than a difference in optimization freedom.

#### 8 Supplementary Note 8: Negative-Binomial Training Objective

Let $L_{s,n} = \sum_{g \in \mathcal{G}} y_{s,n,g}$ be the raw library size at measurement location $n$. The size factor is

$$s_{s,n} = \frac{L_{s,n}}{\frac{1}{N_{\text{train}}} \sum_{(s',n') \in \mathcal{T}} L_{s',n'} + \epsilon},$$

where $\mathcal{T}$ is the training set and $\epsilon = 10^{-8}$. The count-scale mean used in the likelihood is

$$\mu_{s,n,g} = s_{s,n} \bar{\rho}_{s,n,g}.$$

Spatial transcriptomics count data are overdispersed, so Morpho-FM is trained with a gene-wise negative-binomial (NB) objective, consistent with count-modeling practice in single-cell and spatial transcriptomic analysis [17]. With $\theta_g = \exp(\phi_g)$, the log-likelihood contribution for one location and gene is

$$\ell(y_{s,n,g}; \mu_{s,n,g}, \theta_g) = \theta_g \log \frac{\theta_g}{\theta_g + \mu_{s,n,g} + \epsilon} + y_{s,n,g} \log \frac{\mu_{s,n,g} + \epsilon}{\theta_g + \mu_{s,n,g} + \epsilon}$$
$$+ \log \Gamma(y_{s,n,g} + \theta_g + \epsilon) - \log \Gamma(\theta_g + \epsilon) - \log \Gamma(y_{s,n,g} + 1).$$

The optimization objective is the mean negative log-likelihood:

$$\mathcal{L}_{\text{NB}} = -\frac{1}{|\mathcal{T}|\,|\mathcal{G}|} \sum_{(s,n) \in \mathcal{T}} \sum_{g \in \mathcal{G}} \ell(y_{s,n,g}; \mu_{s,n,g}, \theta_g).$$

For large gene panels, the gene set is divided into chunks $\mathcal{C}_1, \ldots, \mathcal{C}_B$. At each training iteration, the chunk order is randomly permuted, and the same morphology-derived representations are supervised against the genes in the current chunk. This implements the same negative-binomial objective while reducing GPU memory requirements.

#### 9 Supplementary Note 9: Dense Decoding and Re-Aggregation

After training, the same adapter and CellFM decoder are applied to every valid tissue-grid location $q = (u, v)$. Let $x_{s,q} = \hat{F}_s(u, v)$. Dense decoding gives

$$\rho_{s,q,g} = \text{softplus}\left(\langle \sigma(W_{\text{dec}} \tilde{e}_g), h_{s,q} \rangle\right).$$

No size factor is applied at unmeasured grid locations, because no library size is observed there. Thus, dense maps are normalized expression-rate fields, not count matrices.

Table 4: Primary training hyperparameters.

| Parameter | Value |
|---|---|
| Optimizer | AdamW |
| Maximum epochs | 100 |
| Weight decay | $10^{-2}$ |
| Adapter and dispersion learning rate | $1 \times 10^{-4}$ |
| Gene embedding and decoder learning rate | $3 \times 10^{-4}$ |
| Early stopping | validation NB loss, patience 5 |
| Minimum improvement | $10^{-4}$ |
| Random seed | 42 |
| Gene chunk size | 256 for large panels; 512 for Xenium |

For quantitative evaluation at the original assay support, dense maps are re-aggregated using the same disk kernel:

$$\rho^{\mathrm{agg}}_{s,n,g} = \frac{1}{|\mathcal{D}(\tilde{r}_s)|} \sum_{(i,j)\in\mathcal{D}(\tilde{r}_s)} \rho_{s,\tilde{p}_{s,n}+(i,j),g}.$$

The correlation between $\rho^{\mathrm{agg}}_{s,n,g}$ and the measured expression $y_{s,n,g}$ is reported as dense-to-measurement re-aggregation consistency. This evaluates whether dense decoding remains coupled to measured expression at locations where measurements exist; it does not provide direct ground truth for unmeasured grid positions.

### 10 Supplementary Note 10: Xenium Breast Reconstruction Analysis

#### 10.1 Four reconstruction settings

The Xenium analysis used two consecutive breast cancer sections, S1/XEN1 and S2/XEN2. After coordinate harmonization and CellFM vocabulary mapping, 306 genes were retained. Four settings were evaluated: S1-to-S1, S2-to-S2, S1-to-S2 and S2-to-S1. Within-section settings test measurement-constrained reconstruction, whereas reciprocal settings test cross-section transfer between consecutive sections.

#### 10.2 Balanced marker panel

In addition to the full 306-gene panel, a fixed 29-gene marker panel was used to summarize tissue-programme recovery across tumour/luminal, epithelial, proliferative, stromal, vascular and immune categories.

Mean per-gene Pearson correlations for the 306-gene observed-location prediction were 0.448 and 0.501 within section, and 0.249 and 0.266 under reciprocal transfer. For the 29-gene marker panel, the corresponding observed-location correlations were 0.651 and 0.708 within section, and 0.396 and 0.427 under transfer. Dense re-aggregation for the marker panel yielded 0.503 and 0.571 within section, and 0.247 and 0.287 under transfer.

Table 5: Xenium marker categories used for balanced panel summaries.

| Category | Genes |
|---|---|
| Tumour/luminal | ERBB2, ESR1, PGR, FOXA1, GATA3 |
| Epithelial | EPCAM, KRT7, KRT8, TACSTD2, CEACAM6 |
| Proliferation | MKI67, TOP2A, CCND1, CENPF |
| Stromal | LUM, POSTN, MMP2, LRRC15, SFRP1 |
| Vascular | PECAM1, VWF, RAMP2, AQP1, CAV1 |
| Immune | PTPRC, CD3D, CD4, CD8A, C1QA |

##### 10.3 Boundary and Region-of-Interest Consistency

Boundary-gradient analysis compared measured and predicted signature profiles as a function of signed distance to the tumour boundary. If $d_n$ is the signed distance of location $n$ to the annotated tumour edge and $S_k$ is the gene set defining signature $k$, the signature score at location $n$ was computed as an average over the retained signature genes:

$$A_{n,k} = \frac{1}{|S_k|} \sum_{g \in S_k} \tilde{y}_{n,g},$$

where $\tilde{y}_{n,g}$ denotes normalized measured or predicted expression. Scores were averaged within distance bins and compared between measured and predicted profiles. Signature-wise boundary correlations were 0.954–0.985 within section and 0.871–0.970 under cross-section transfer.

Region-of-interest (ROI) enrichment analysis used curated tissue regions. For ROI $R$ and signature $S_k$, enrichment was computed from the mean signature score inside the ROI relative to the section-level background. Across 80 ROI–signature pairs, measured-versus-predicted enrichment correlations were 0.889 within section and 0.770 under transfer.

##### 10.4 Latent-domain clustering

For Xenium domain analysis, the 1,536-dimensional CellFM-derived latent representation before gene-specific readout was extracted at valid locations. Principal component analysis (PCA) was applied before MiniBatch K-means clustering. The default resolution was $k = 15$, and robustness was assessed for $k = 10, 15, 20$ across five random seeds. Agreement with curated coarse tissue annotations was quantified by normalized mutual information (NMI) and adjusted Rand index (ARI). At $k = 10$, Morpho-FM latent clustering reached NMI/ARI $0.396 \pm 0.004/0.285 \pm 0.021$; at $k = 15$, $0.376 \pm 0.006/0.227 \pm 0.026$; and at $k = 20$, $0.362 \pm 0.004/0.175 \pm 0.009$. Raw morphology clustering was evaluated under the same PCA and clustering procedure as a control.

### 11 Supplementary Note 11: Evaluation Metrics

#### 11.1 Per-gene Pearson correlation

For gene $g$, prediction quality was measured by Pearson correlation across held-out measurement locations:

$$r_g = \frac{\sum_n (\hat{y}_{n,g} - \bar{\hat{y}}_g)(y_{n,g} - \bar{y}_g)}{\sqrt{\sum_n (\hat{y}_{n,g} - \bar{\hat{y}}_g)^2}\sqrt{\sum_n (y_{n,g} - \bar{y}_g)^2}}.$$

Genes with undefined correlation because of zero variance were excluded from mean and median summaries using finite-value aggregation.

#### 11.2 Highly Variable Gene Recovery

Highly variable gene (HVG) analyses were post hoc and followed the common use of variable-feature selection to summarize strong expression heterogeneity [16, 17]. For target section $s$, the measured HVG set $\mathcal{H}_K^{\text{true}}$ was defined from the measured expression matrix, and the predicted HVG set $\mathcal{H}_K^{\text{pred}}$ was defined from the predicted matrix. Recall@$K$ was

$$\text{Recall@}K = \frac{|\mathcal{H}_K^{\text{true}} \cap \mathcal{H}_K^{\text{pred}}|}{K}.$$

Prediction quality for measured HVGs was summarized by the mean of $r_g$ over $\mathcal{H}_{1000}^{\text{true}}$.

#### 11.3 Matrix-level and domain-level metrics

For external kidney transfer, global expression correlation was computed by vectorizing the full location-by-gene matrix. Spot-profile correlation was computed per location and then averaged:

$$r_n^{\text{spot}} = \text{corr}\left(\hat{y}_{n,\mathcal{G}}, y_{n,\mathcal{G}}\right).$$

For domain analyses, NMI and ARI were used to compare inferred clusters with curated annotations where such annotations were available. Boundary and ROI analyses were treated as biological consistency checks and were not used for training or checkpoint selection.

### 12 Supplementary Note 12: Reproducibility and Auditability

All benchmark comparisons were recomputed from saved prediction and reference matrices after output alignment. For each run, the retained gene list, measurement-location order, split definition and metric summary were preserved. This makes the benchmark auditable at the level of the evaluated matrices rather than only at the level of final plots. The released code includes Morpho-FM training, inference, dense re-aggregation and benchmark evaluation workflows.
