## Supplementary Figure for "Morpho-FM: spatial molecular reconstruction from routine H&E histology using transcriptomic foundation-model priors"

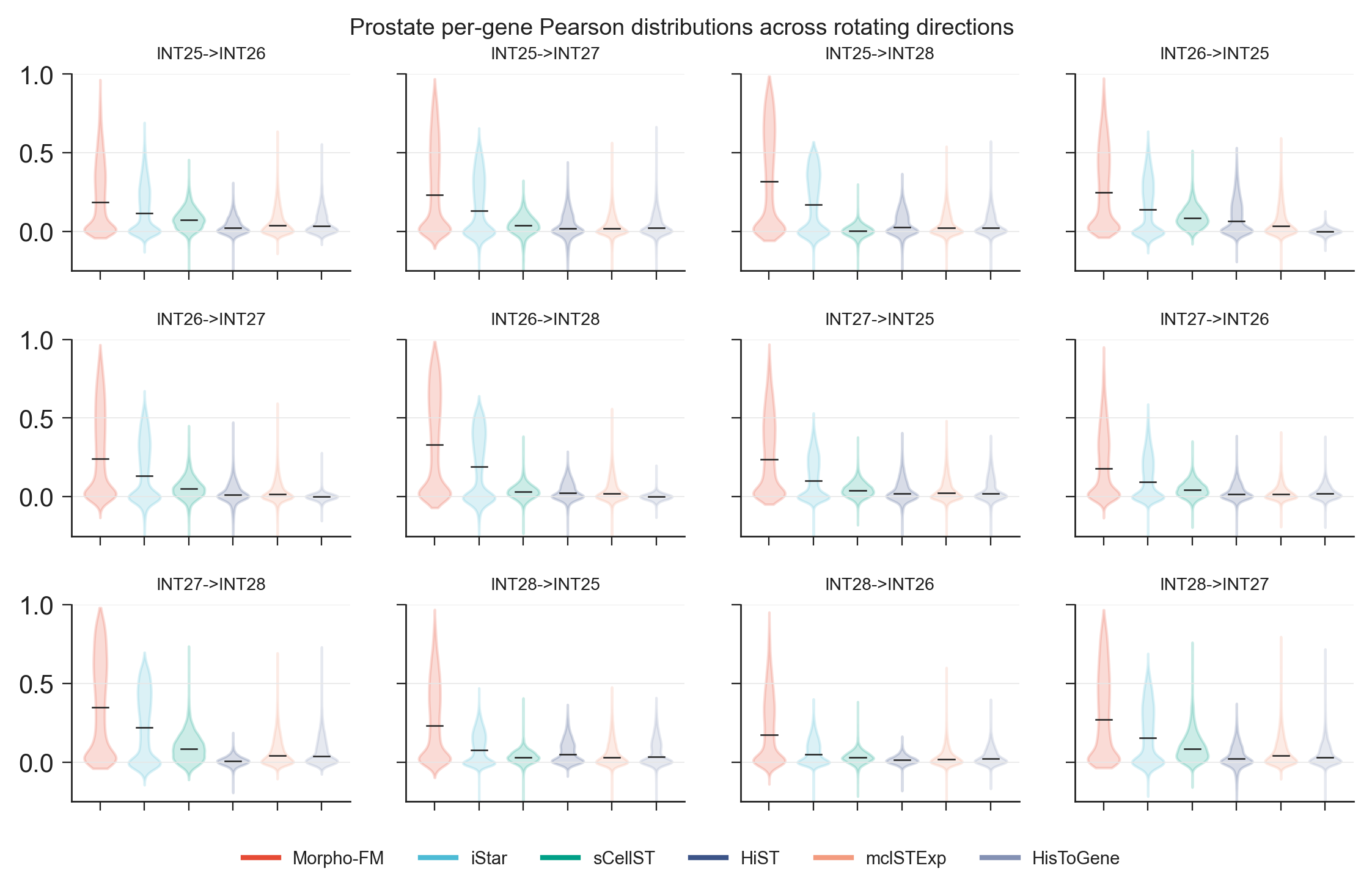

**Supplementary Fig. 1 | Per-gene prediction performance across the rotating single-slide cross-section prostate benchmark.** Violin plots show the distribution of per-gene Pearson correlation coefficients across all 12 train–test directions in the HEST24 prostate cohort. In each direction, one prostate section was used as the only training slide and a different section was held out for testing, as indicated by the title above each panel. Prediction performance was evaluated on the shared prostate gene panel for Morpho-FM, iStar, sCellST, HiST, mclSTExp and HisToGene. Across the rotating directions, Morpho-FM consistently produced a right-shifted per-gene correlation distribution and higher central performance than the competing methods, demonstrating that its cross-section generalization advantage was maintained across different training and held-out test sections rather than being driven by a single favorable comparison.

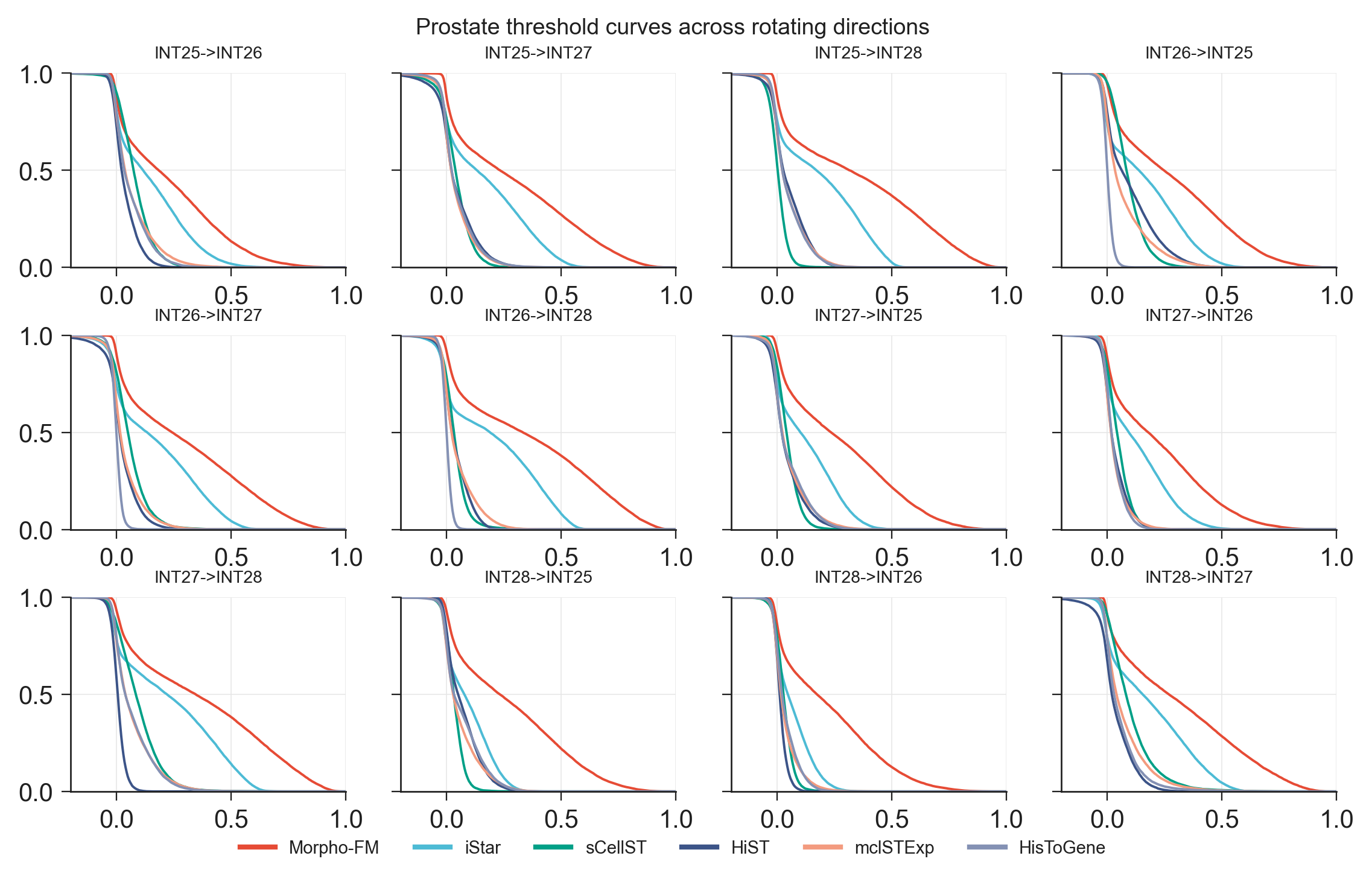

**Supplementary Fig. 2 | Threshold-based gene recovery across the rotating single-slide cross-section prostate benchmark.** Threshold curves show the fraction of genes exceeding increasing Pearson correlation thresholds across all 12 train–test directions in the HEST24 prostate cohort. In each direction, one prostate section was used as the only training slide and a different section was held out for testing, as indicated above each panel. Prediction performance was evaluated on the shared prostate gene panel for Morpho-FM, iStar, sCellST, HiST, mclSTExp and HisToGene. Across the rotating directions, Morpho-FM retained a larger fraction of genes above progressively more stringent Pearson thresholds than the competing methods, indicating that its cross-section advantage reflected broad improvement across many genes rather than isolated gains in a small subset of highly predictable genes.

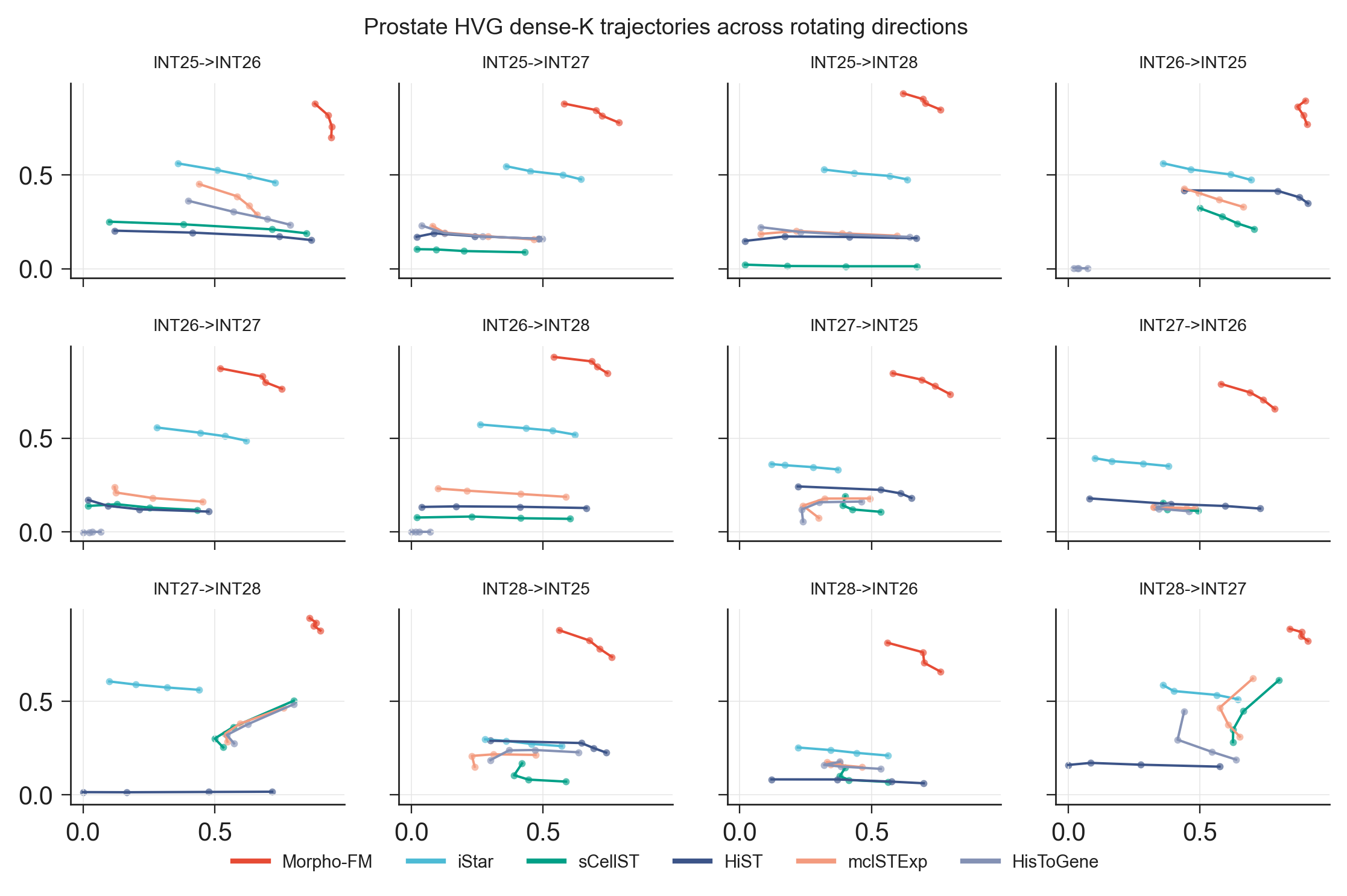

**Supplementary Fig. 3 | HVG recovery and prediction quality across the rotating single-slide cross-section prostate benchmark.** HVG-focused top-K trajectories across all 12 train–test directions in the HEST24 prostate cohort. In each direction, one prostate section was used as the only training slide and a different section was held out for testing, as indicated above each panel. Each trajectory summarizes performance across increasing top-K cutoffs, comparing HVG overlap with the observed data (Recall@K; x axis) and the mean Pearson correlation of the predicted HVG set (y axis). Across the rotating directions, Morpho-FM generally occupied the upper region of the plots, achieving substantially higher prediction accuracy for HVG-enriched gene sets than iStar, sCellST, HiST, mclSTExp and HisToGene. These results show that Morpho-FM’s cross-section advantage was preserved for biologically variable genes rather than being limited to average performance across the full gene panel.

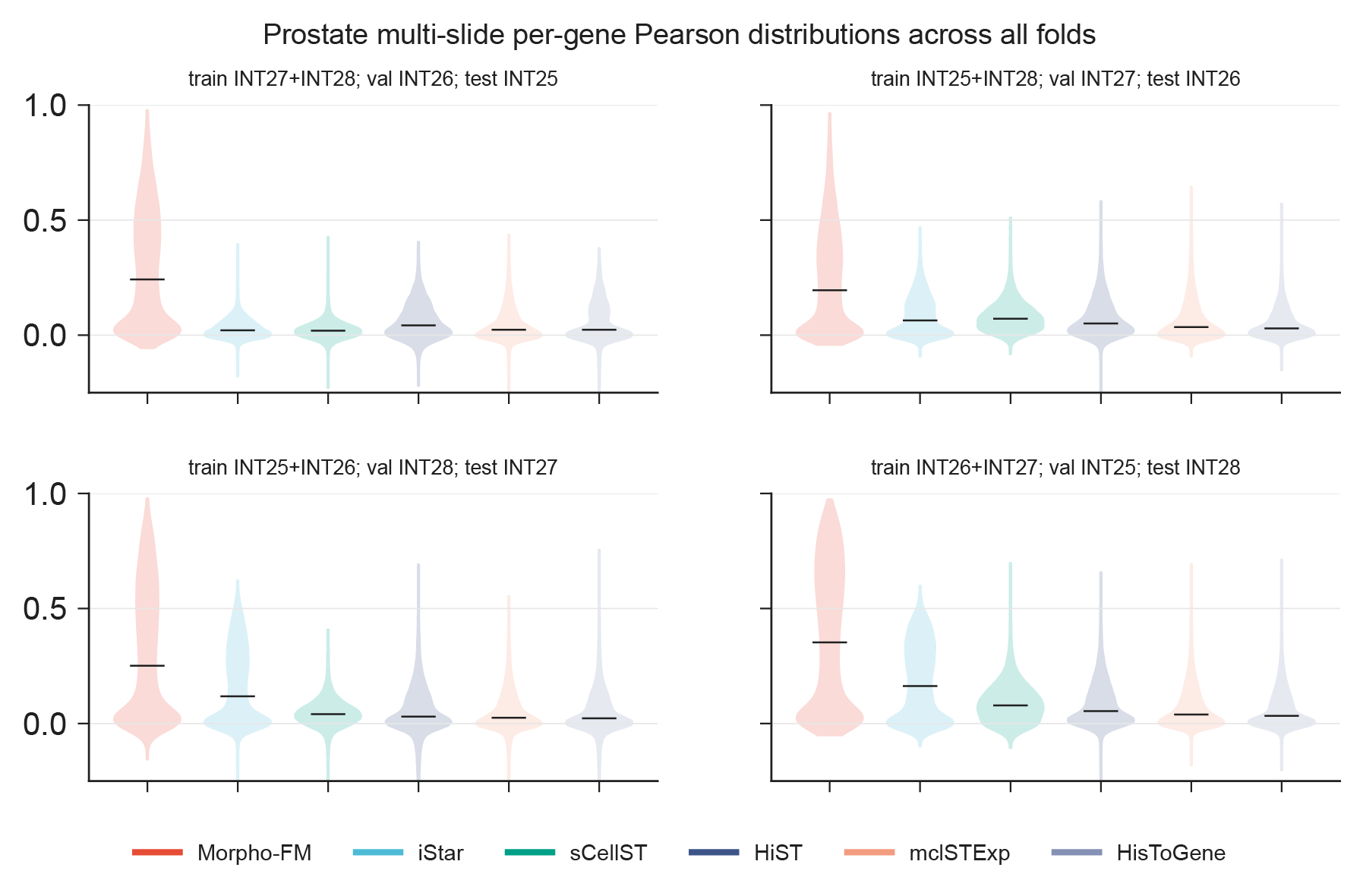

**Supplementary Fig. 4 | Per-gene prediction performance across all folds in the multi-slide prostate benchmark.** Violin plots show the distribution of per-gene Pearson correlation coefficients across the four rotating folds of the multi-slide prostate benchmark. In each fold, two prostate sections were used for training, a third section was used for validation and the remaining section was held out for testing, as indicated above each panel. Prediction performance was evaluated on the shared prostate gene panel for Morpho-FM, iStar, sCellST, HiST, mclSTExp and HisToGene. Across all held-out test sections, Morpho-FM produced consistently right-shifted per-gene correlation distributions and higher central performance than the competing methods, showing that its cross-section generalization advantage was maintained in the more data-rich multi-slide setting rather than being restricted to the single-slide benchmark.

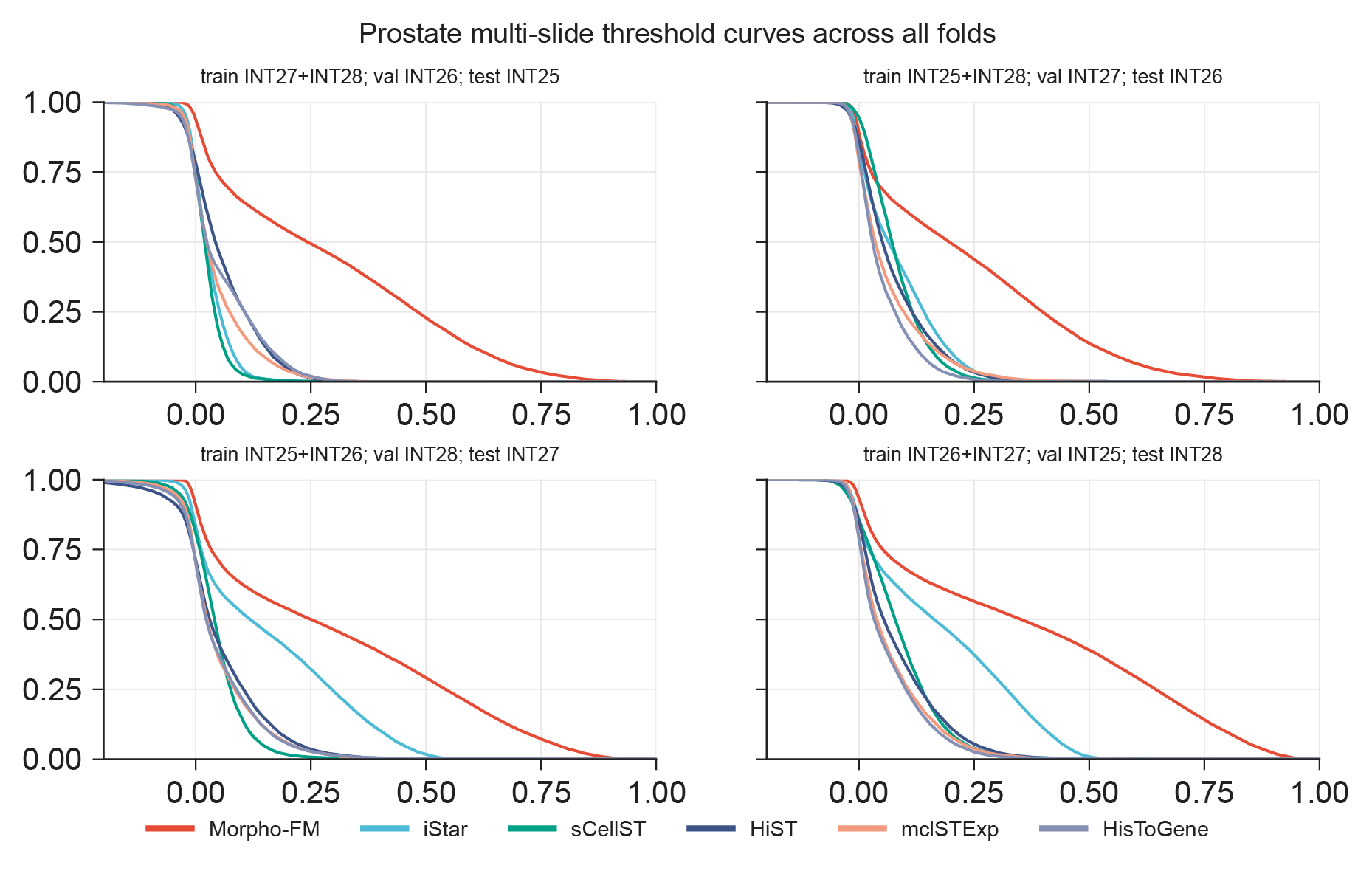

**Supplementary Fig. 5 | Threshold-based gene recovery across all folds in the multi-slide prostate benchmark.** Threshold curves show the fraction of genes exceeding increasing Pearson correlation thresholds across the four rotating folds of the multi-slide prostate benchmark. In each fold, two prostate sections were used for training, a third section was used for validation and the remaining section was held out for testing, as indicated above each panel. Prediction performance was evaluated on the shared prostate gene panel for Morpho-FM, iStar, sCellST, HiST, mclSTExp and HisToGene. Across all held-out test sections, Morpho-FM retained a larger fraction of genes above progressively more stringent Pearson thresholds than the competing methods, indicating that its advantage in the multi-slide prostate benchmark reflected broadly improved gene-level prediction rather than isolated gains in a small subset of genes.

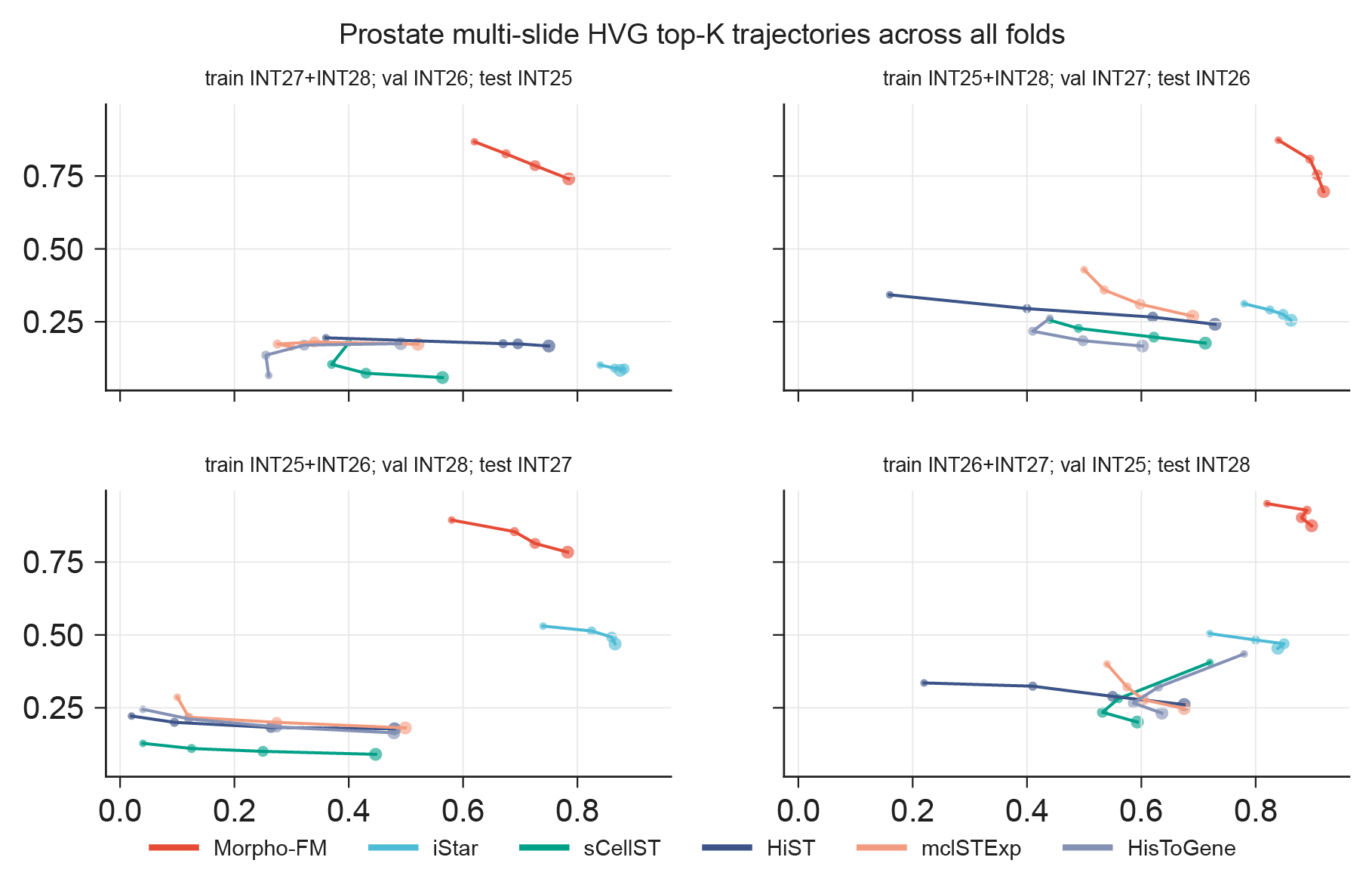

**Supplementary Fig. 6 |** **HVG recovery and prediction quality across all folds in the multi-slide prostate benchmark.** HVG-focused top-K trajectories across the four rotating folds of the multi-slide prostate benchmark. In each fold, two prostate sections were used for training, a third section was used for validation and the remaining section was held out for testing, as indicated above each panel. Each trajectory summarizes performance across increasing top-K cutoffs by comparing HVG overlap with the observed data (Recall@K; x axis) and the mean Pearson correlation of the predicted HVG set (y axis). Across all held-out test sections, Morpho-FM achieved the strongest HVG-focused performance, occupying the upper-right region of the plots and showing higher prediction accuracy for biologically variable genes than iStar, sCellST, HiST, mclSTExp and HisToGene. These results indicate that Morpho-FM’s advantage in the multi-slide prostate benchmark was preserved for highly variable genes rather than being limited to average performance across the full gene panel.

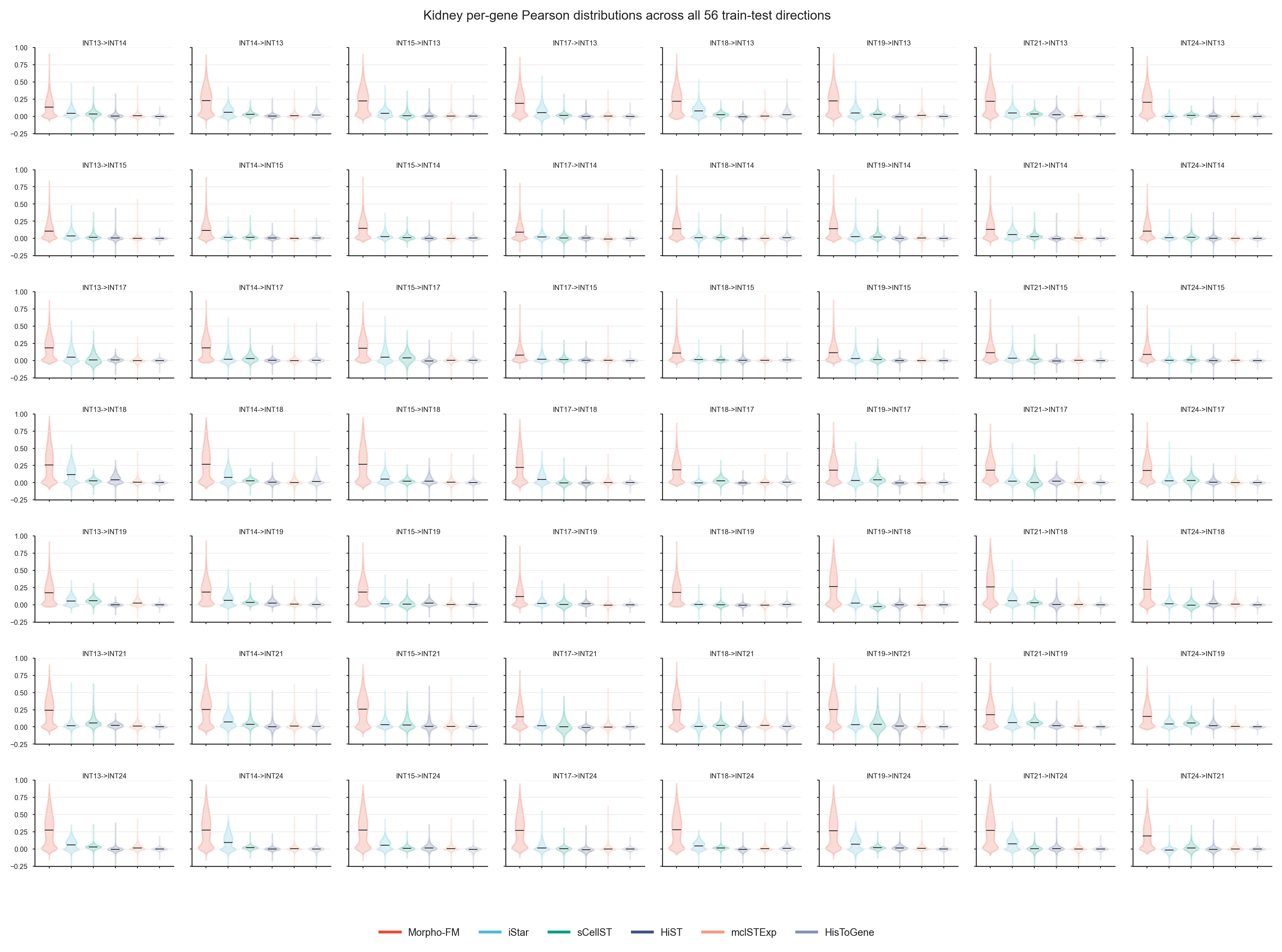

**Supplementary Fig. 7 |** **Per-gene prediction performance across the rotating single-slide cross-section kidney cancer benchmark.** Violin plots show the distribution of per-gene Pearson correlation coefficients across all 56 train–test directions in the kidney cancer cohort. In each direction, one kidney cancer section was used as the only training slide and a different section was held out for testing, as indicated above each panel. Prediction performance was evaluated on the shared kidney gene panel for Morpho-FM, iStar, sCellST, HiST, mclSTExp and HisToGene. Across the rotating directions, Morpho-FM consistently produced right-shifted per-gene correlation distributions and higher central performance than the competing methods, demonstrating stable single-slide cross-section generalization across different training and held-out kidney cancer sections.

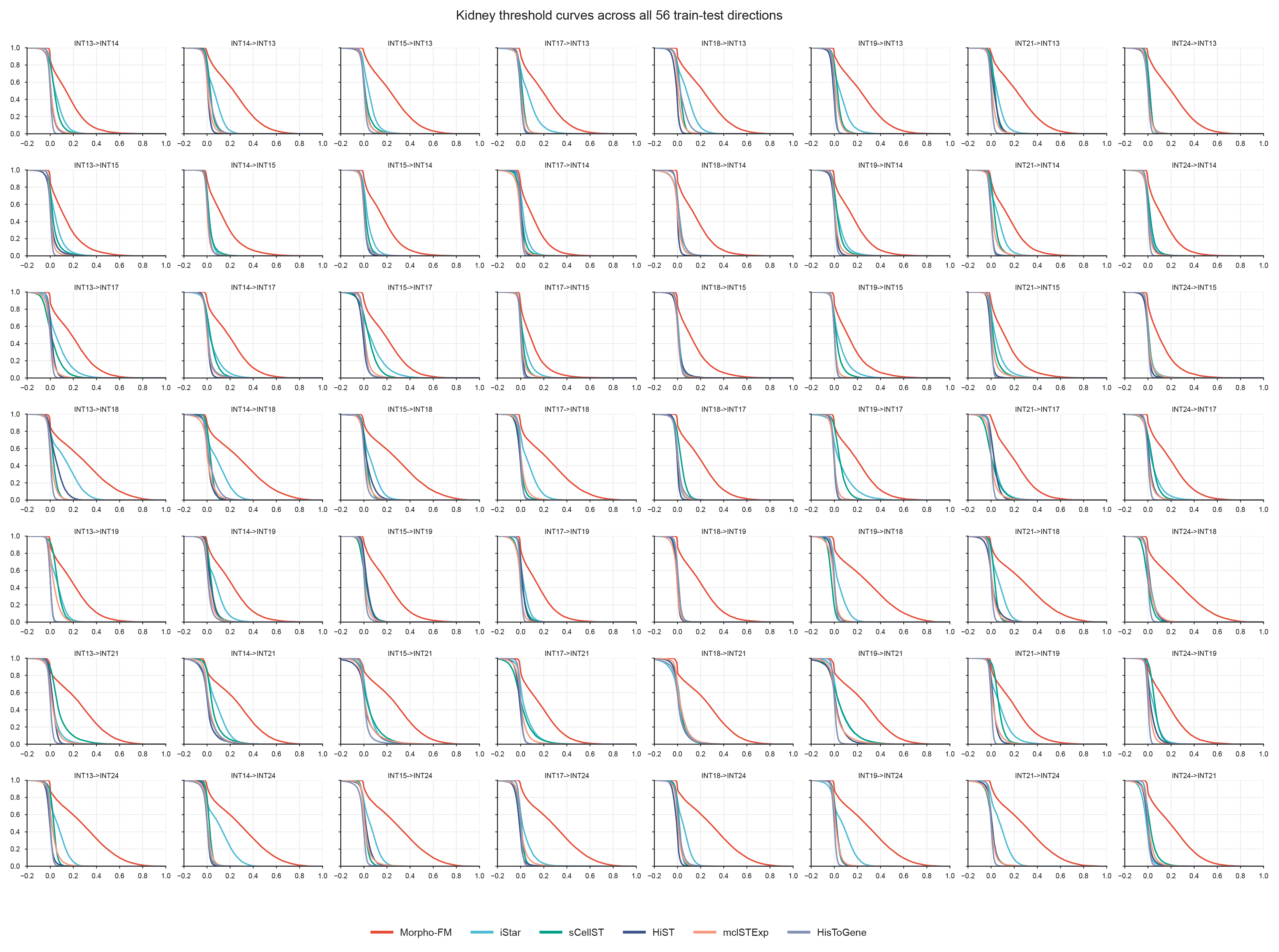

**Supplementary Fig. 8 | Threshold-based gene recovery across the rotating single-slide cross-section kidney cancer benchmark.** Threshold curves show the fraction of genes exceeding increasing Pearson correlation thresholds across all 56 train–test directions in the kidney cancer cohort. In each direction, one kidney cancer section was used as the only training slide and a different section was held out for testing, as indicated above each panel. Prediction performance was evaluated on the shared kidney gene panel for Morpho-FM, iStar, sCellST, HiST, mclSTExp and HisToGene. Across the rotating directions, Morpho-FM retained a larger fraction of genes above progressively more stringent Pearson thresholds than the competing methods, indicating that its single-slide cross-section advantage reflected broad improvement across many genes rather than isolated gains in a small subset of highly predictable genes.

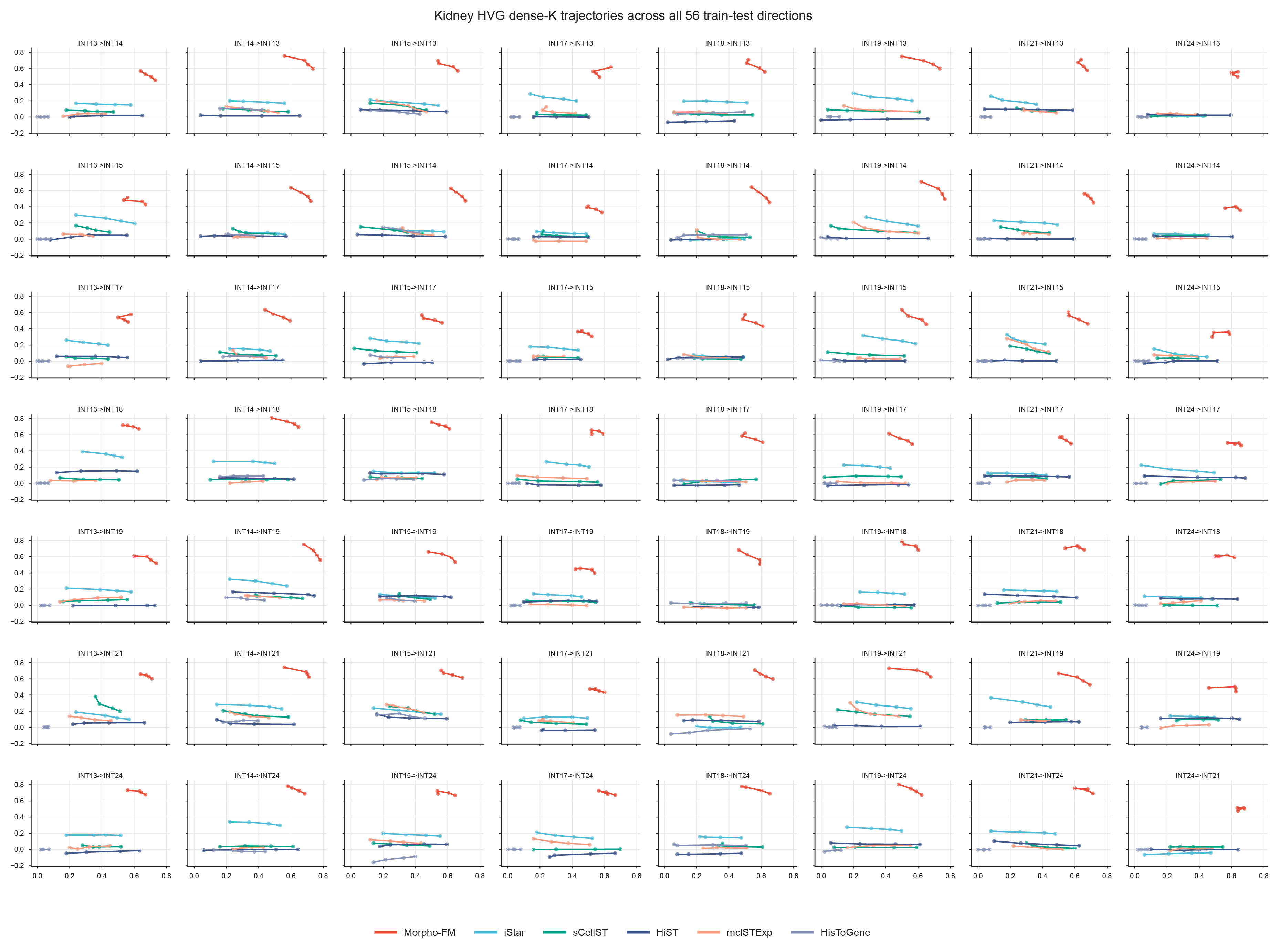

**Supplementary Fig. 9 | HVG recovery and prediction quality across the rotating single-slide cross-section kidney cancer benchmark.** HVG-focused top-K trajectories across all 56 train–test directions in the kidney cancer cohort. In each direction, one kidney cancer section was used as the only training slide and a different section was held out for testing, as indicated above each panel. Each trajectory summarizes performance across increasing top-K cutoffs by comparing HVG overlap with the observed data (Recall@K; x axis) and the mean Pearson correlation of the predicted top-K gene set (y axis). Across the rotating directions, Morpho-FM generally occupied the upper region of the plots and achieved higher prediction accuracy for HVG-enriched gene sets than iStar, sCellST, HiST, mclSTExp and HisToGene. These results indicate that Morpho-FM’s single-slide cross-section advantage in the kidney cancer cohort was preserved for biologically variable genes rather than being limited to average performance across the full gene panel.

**Supplementary Fig. 10 | Additional marker-gene spatial comparisons after external transfer to ccRCC sections. a,** Measured and predicted spatial expression maps for *AQP1*, *VCAM1*, *TFEB* and *SDHB* in external ccRCC section 1. The measured expression map is shown in the top row and the Morpho-FM prediction is shown in the bottom row. Pearson correlation coefficients between predicted and measured expression across measurement locations are shown above each gene. **b,** Measured and predicted spatial expression maps for *SMARCB1*, *TFE3*, *TFEB* and *SDHB* in external ccRCC section 2, shown as in **a**. Predictions were generated using the kidney-trained Morpho-FM model without additional fitting or target-section-specific tuning. Colour scales indicate relative expression from low to high within each gene.

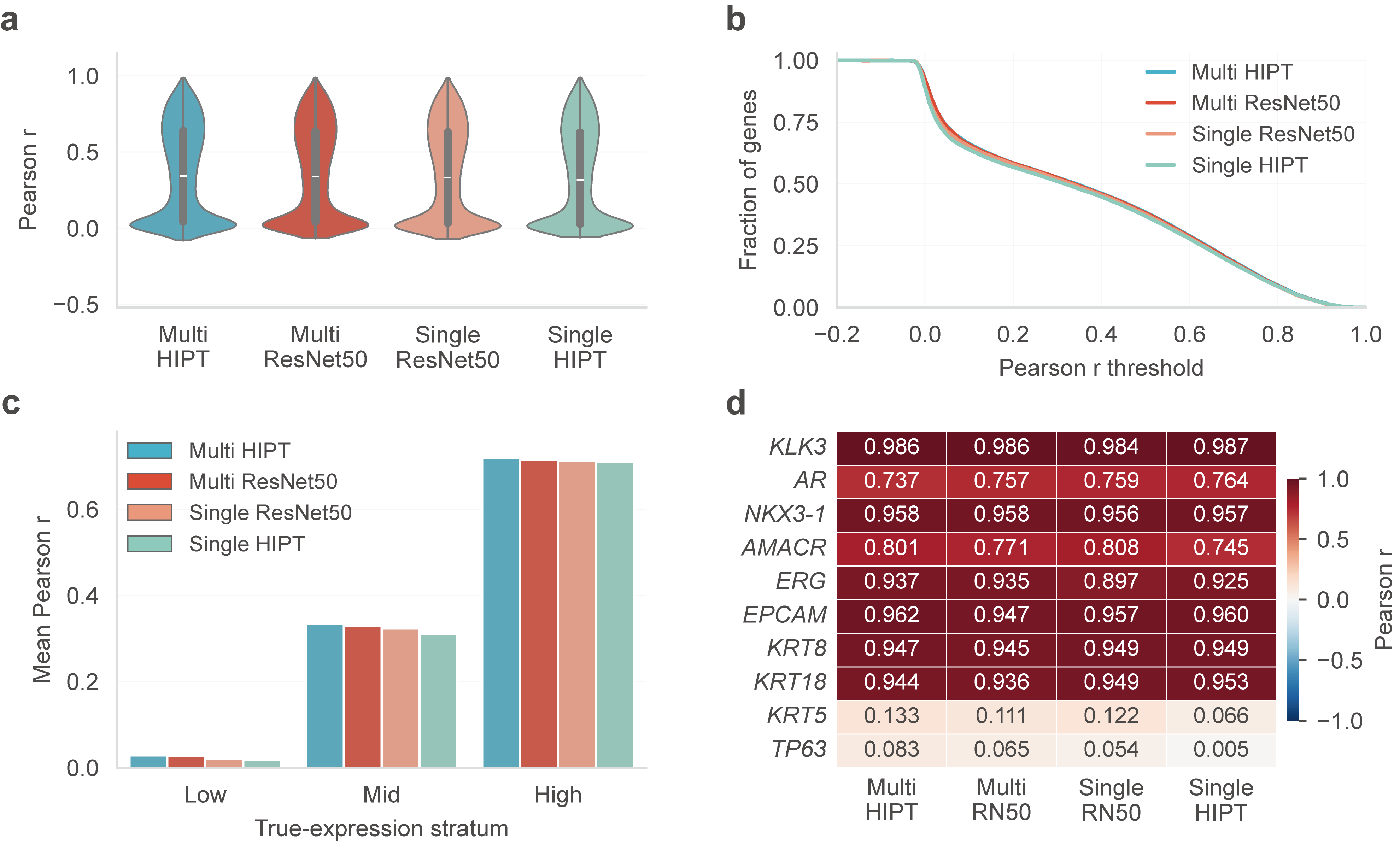

**Supplementary Fig. 11 | Comparison of HIPT and ResNet-50 visual features in Morpho-FM.**
**a,** Per-gene Pearson correlation distributions for Morpho-FM models using HIPT features or ImageNet-pretrained ResNet-50 features under matched multi-slide and single-slide prostate training settings. **b,** Fraction of genes with Pearson correlation above each threshold for the four visual-encoder variants shown in **a**. **c,** Mean per-gene Pearson correlation stratified by measured expression level. Genes were grouped into low-, mid- and high-expression strata based on their true expression levels, and performance is shown for each visual-encoder variant. **d,** Gene-level Pearson correlations for a fixed prostate marker panel across the four visual-encoder variants. Heatmap values indicate Pearson correlation coefficients between predicted and measured expression for each marker gene.

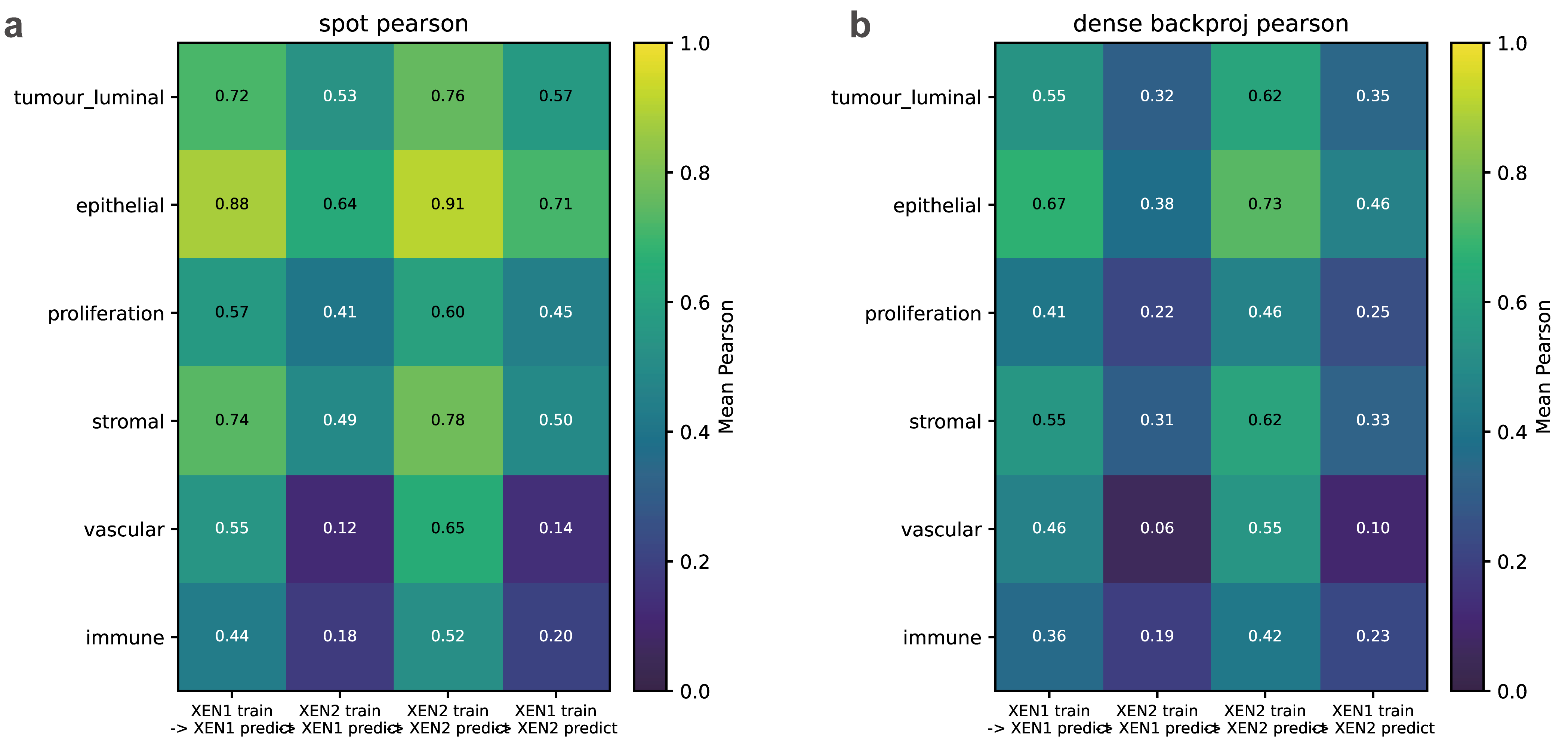

**Supplementary Fig. 12 | Four-setting Xenium prediction distributions and marker-category summaries. a,** Per-gene Pearson correlation distributions at the observed Xenium measurement locations across the four within-section and reciprocal section-transfer settings. Results are shown for all 306 retained genes and for the fixed 29-gene breast marker panel. **b,** Dense-to-measurement re-aggregation performance for the same 29-gene marker panel. Marker-category summaries show performance across tumour/luminal, epithelial, proliferation, stromal, vascular and immune programmes, supporting the main-text observation that epithelial markers were more reproducible across sections whereas vascular and immune markers showed larger cross-section loss.

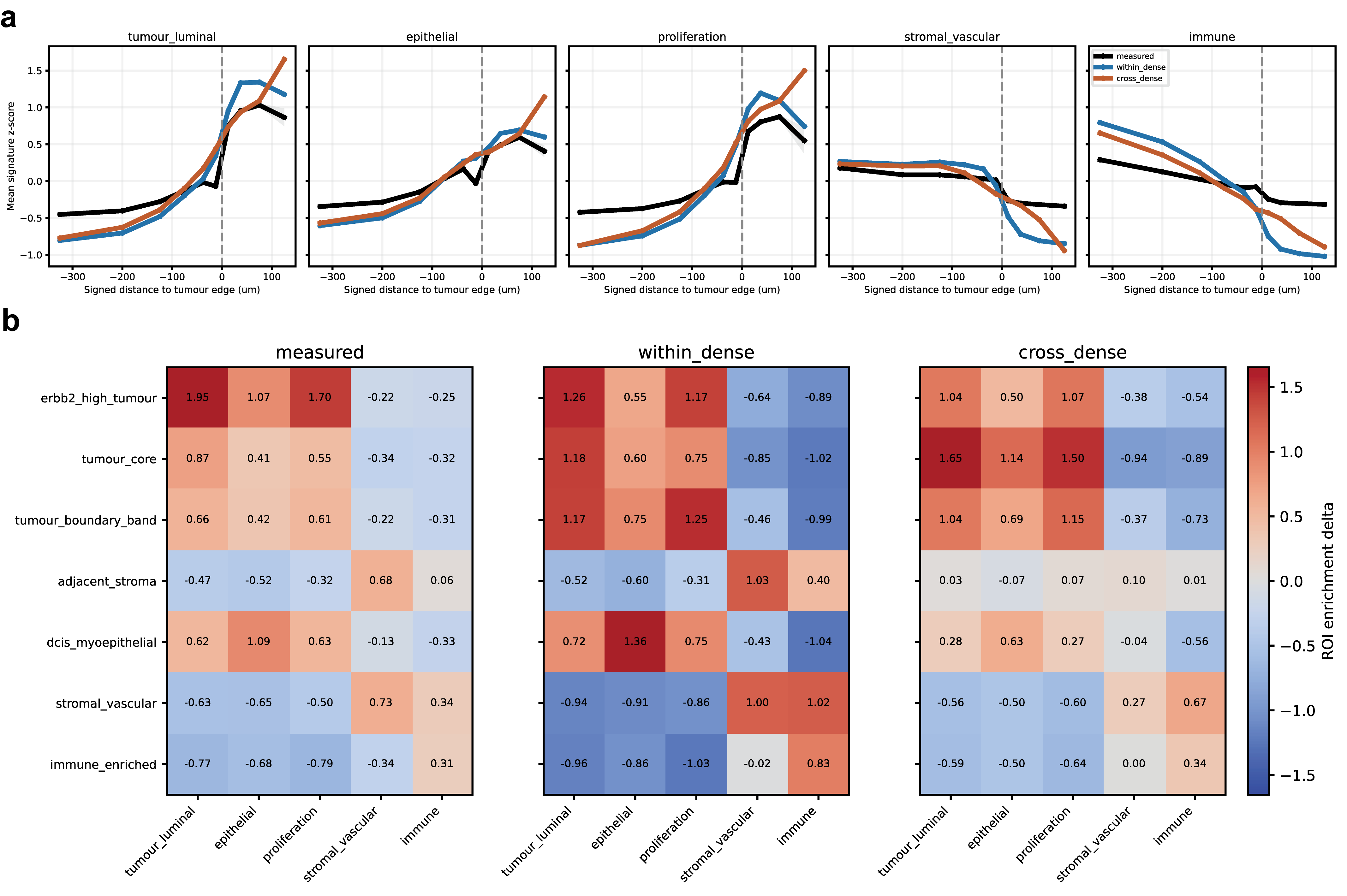

**Supplementary Fig. 13 | Boundary-gradient and ROI-level agreement analyses in Xenium S1.**
**a,** Signature-level boundary-gradient profiles comparing measured Xenium expression with within-section dense reconstruction and reciprocal cross-section dense prediction across signed distance bins from the tumour boundary. Signatures span tumour/luminal, epithelial, proliferation, stromal/vascular and immune programmes. **b,** ROI-level signature enrichment comparison across biologically defined regions, including ERBB2-high tumour, tumour core, tumour boundary, adjacent stroma, DCIS/myoepithelial, stromal/vascular and immune-enriched regions. These analyses test whether dense predictions preserve spatial expression trends across tissue boundaries and regions of interest beyond pointwise gene-level correlation.

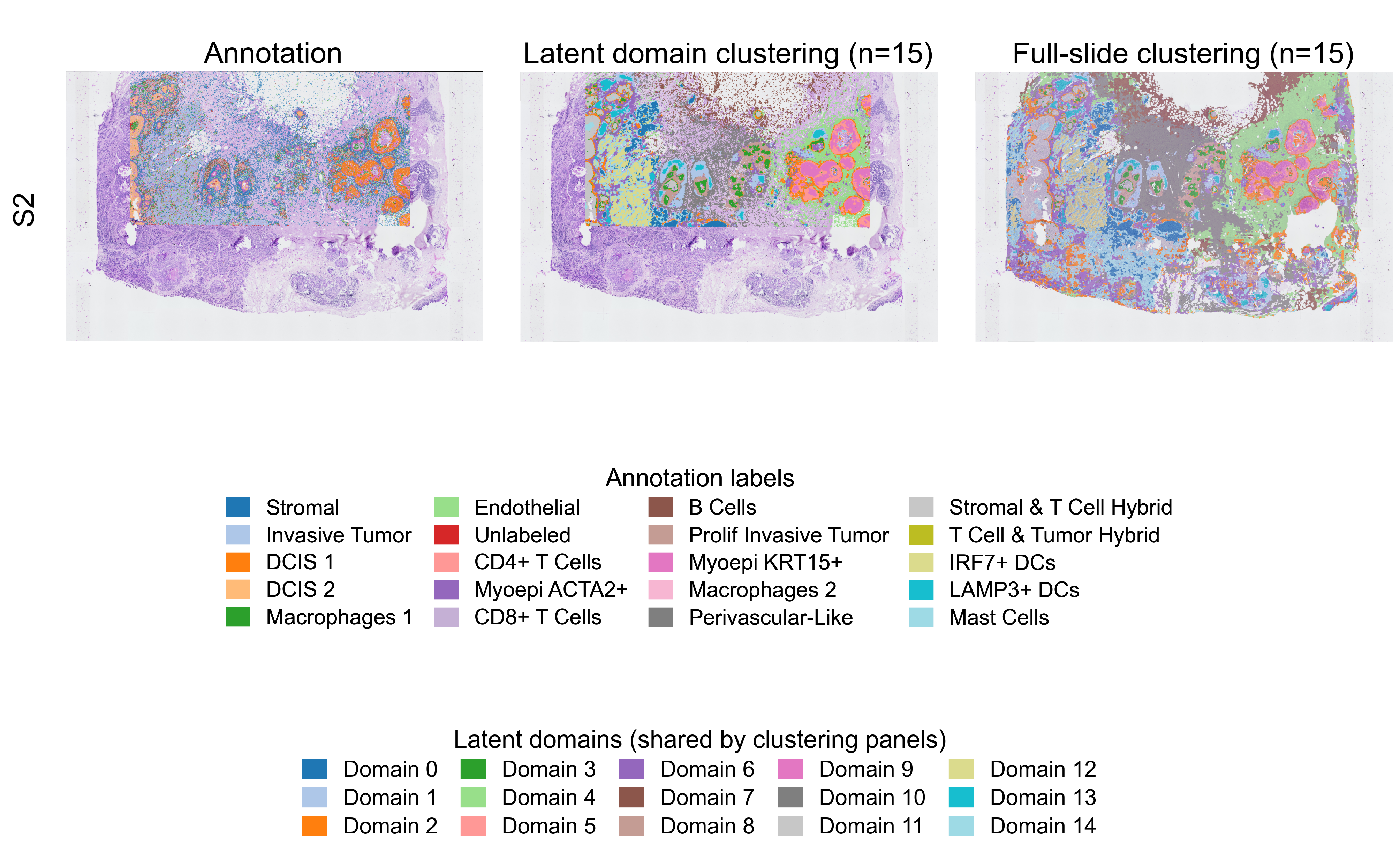

**Supplementary Fig. 14 | Annotation and latent-domain structure in Xenium section 2.** Cell-level annotation overlay, Morpho-FM latent domain clustering at observed Xenium measurement locations, and full-slide latent domain clustering for Xenium section 2 (S2). Clustering was performed with k = 15. Annotation labels are shown with a shared colour key, and the two clustering panels use a common latent-domain colour key.

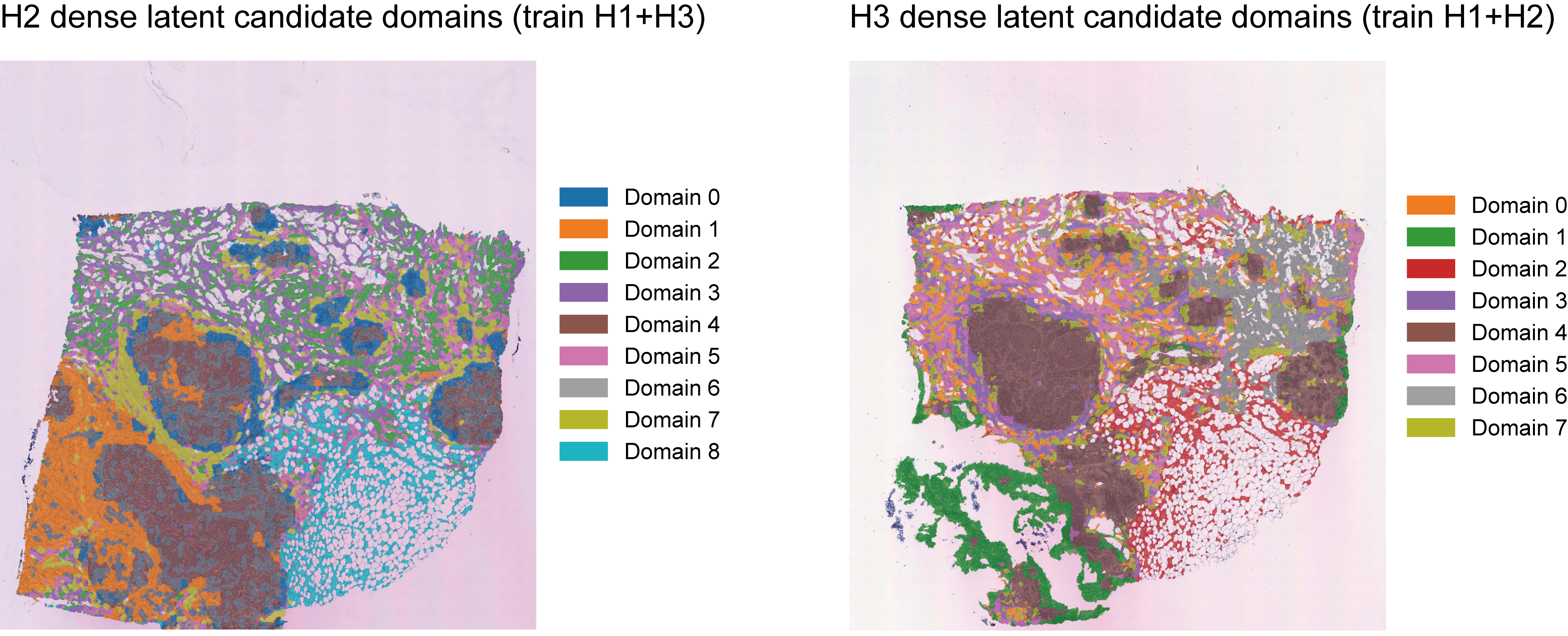

**Supplementary Fig. 15 |** **Dense latent candidate domains in held-out HER2ST H2 and H3 sections.** Dense latent-domain maps are shown for the unlabeled HER2ST patient H sections H2 and H3 under leave-one-section-out transfer. For H2, Morpho-FM was trained on H1 and H3 and applied to the held-out H2 section; for H3, the model was trained on H1 and H2 and applied to the held-out H3 section. Domains were obtained by clustering dense CellFM-derived latent embeddings before gene-specific readout and are overlaid on the corresponding H&E images. Colours denote unsupervised candidate domains within each section; domain numbers are section-specific and should not be interpreted as manually validated tissue classes. These maps were used as exploratory spatial-domain outputs to support marker-guided inspection in unlabeled HER2ST sections.
